# Comprehensive study on ferredoxin isoforms in the cyanobacterium Synechocystis sp. PCC 6803

**DOI:** 10.64898/2026.04.08.717189

**Authors:** Marko Boehm, Drazenka Svedruzic, Carolyn E. Lubner, Jens Appel, David W. Mulder, Effie C. Kisgeropoulos, Vanessa Hueren, Katharina Spengler, Vivek S. Bharadwaj, Zhanjun Guo, Anastasia E. Ledinina, Darja Deobald, Lorenz Adrian, Paul W. King, Kirstin Gutekunst

## Abstract

Ferredoxins are central to cellular metabolism by mediating electron flow in energy conversion reactions. The focus of this study was to systematically examine twelve ferredoxin and ferredoxin-like proteins from *Synechocystis* sp. PCC 6803 to identify their properties, activities, and functions in electron transfer. Using electron paramagnetic resonance spectroscopy, we detected cluster types consistent with major ferredoxin families including plant-type [2Fe-2S], adrenodoxin, thioredoxin, and bacterial-type [4Fe– 4S] ferredoxins. In addition, we found that the ssr3184 ferredoxin-like protein exchanged between a [3Fe–4S] or a [4Fe–4S] cluster, pointing to a possible functional change in response to changes in oxygen or cellular redox poise. Electrochemical measurements demonstrated that these ferredoxins constitute a broad potential window, from −243 mV to −520 mV vs SHE. Investigations on their capacity to support electron-transfer focused on reactions with two major redox hubs: Photosystem I and pyruvate:ferredoxin oxidoreductase and included testing of binding interactions with nitrite reductase. Expression profiling under multiple environmental conditions was also used to predict function and revealed distinct regulatory patterns. Collectively, these findings identified a group of core ferredoxins that directly support photosynthetic electron transfer, and more specialized ones that may serve other functions. In summary, *Synechocystis* utilizes a suite of ferredoxins to maintain cellular redox homeostasis under dynamic environmental conditions.

## Introduction

Every living cell depends on a finely regulated distribution of electrons between a multitude of enzymatic reactions. This complex distribution is the basis for a functioning metabolism and redox homeostasis and is required to adapt to changes in environmental conditions. Ferredoxins (Fdx) play a central role in these distribution and adaptation processes. They are small redox proteins with [2Fe-2S], [3Fe-4S], and [4Fe-4S] clusters, in which the irons exist in different oxidation states to shuttle electrons. Fdxs are ubiquitous in all domains of life and among the oldest proteins that emerged during the origin of life on Earth (1). [4Fe-4S] clusters evolved in an anaerobic environment. The emergence of oxygen-tolerant [2Fe–2S] cluster Fdxs marked a major evolutionary turning point, coinciding with the advent of oxygenic photosynthesis (2, 3).

Despite their small size, Fdxs differ in cluster composition, redox potential, and interaction partners, allowing them to participate in a wide range of electron transfer reactions (3-6). In photosynthetic organisms, Fdxs link the photosynthetic electron transport chain to various downstream enzymes involved in cellular metabolism and adaptation to environmental conditions. Fdx diversity reflects an evolutionary specialization in which individual proteins evolved to support diverse reactions, suggesting there has been an evolutionary pressure to select properties for fine-tuning redox flow within specific metabolic contexts and more than simply serving as interchangeable electron carriers.

The model cyanobacterium *Synechocystis* sp. PCC 6803 (hereafter *Synechocystis*) encodes a diverse set of iron-sulfur (Fe-S) proteins (Gene ID, Table 1), some with confirmed and others with putative Fdx or Fdx-like functions (for simplicity we will refer to all as “Fdxs” throughout text). Fdxs 1-4 have been classified as plant-type Fdxs that typically function in photosynthetic electron transfer. Fdx1-Fdx4 have redox midpoint values (*E*_m_) that range from -243 to -460 mV vs SHE (3, 5), suggesting functional diversity. Fdx1, is the most studied Fdx and has the primary function in photosynthetic electron transfer from PSI to Fdx–NADP^+^reductase (FNR) facilitating NADPH use in the Calvin– Benson–Bassham (CBB) cycle (6). Fdx1 also functions in electron transfer to nitrite or nitrate reductases (7), sulfite reductase (8), Fdx-dependent GOGAT (9, 10), [NiFe]-hydrogenase complexes HoxEFUYH (11) and HoxEFU (12, 13). In addition to NAD(P)+, (14) it may donate electrons to flavodiiron proteins Flv1 and Flv3 (15, 16). Fdx2, is an essential Fdx with a high (positive) redox potential (*E*_m_ = -243 mV) (17). Previous studies suggested that Fdx2 is not involved in photosynthetic electron transfer. Under low-iron conditions, cells with lowered Fdx2 levels fail to accumulate the light-harvesting protein IsiA, suggesting that Fdx2 helps coordinate redox signaling with iron homeostasis (17). Fdx3 expression is light-dependent with deletion mutants being impaired in photomixotrophic growth, with pyruvate:ferredoxin oxidoreductase (PFOR) as its electron donor (18). Fdx4 is a low potential ferredoxin, *E*_m_ = -460 mV that is induced under CO_2_, nitrate, or phosphate limitation (19), as well as oxidative stress. Fdx4 also supports NAD^+^ reduction by HoxEFU (12). Fdx5 and Fdx6, are non-essential [2Fe–2S] proteins with unconfirmed function. Fdx5 is co-transcribed with Fdx4, suggesting coordinated regulation under nutrient limitation, while Fdx6 is upregulated under blue light and phosphate deficiency (20).

**Table 1.**
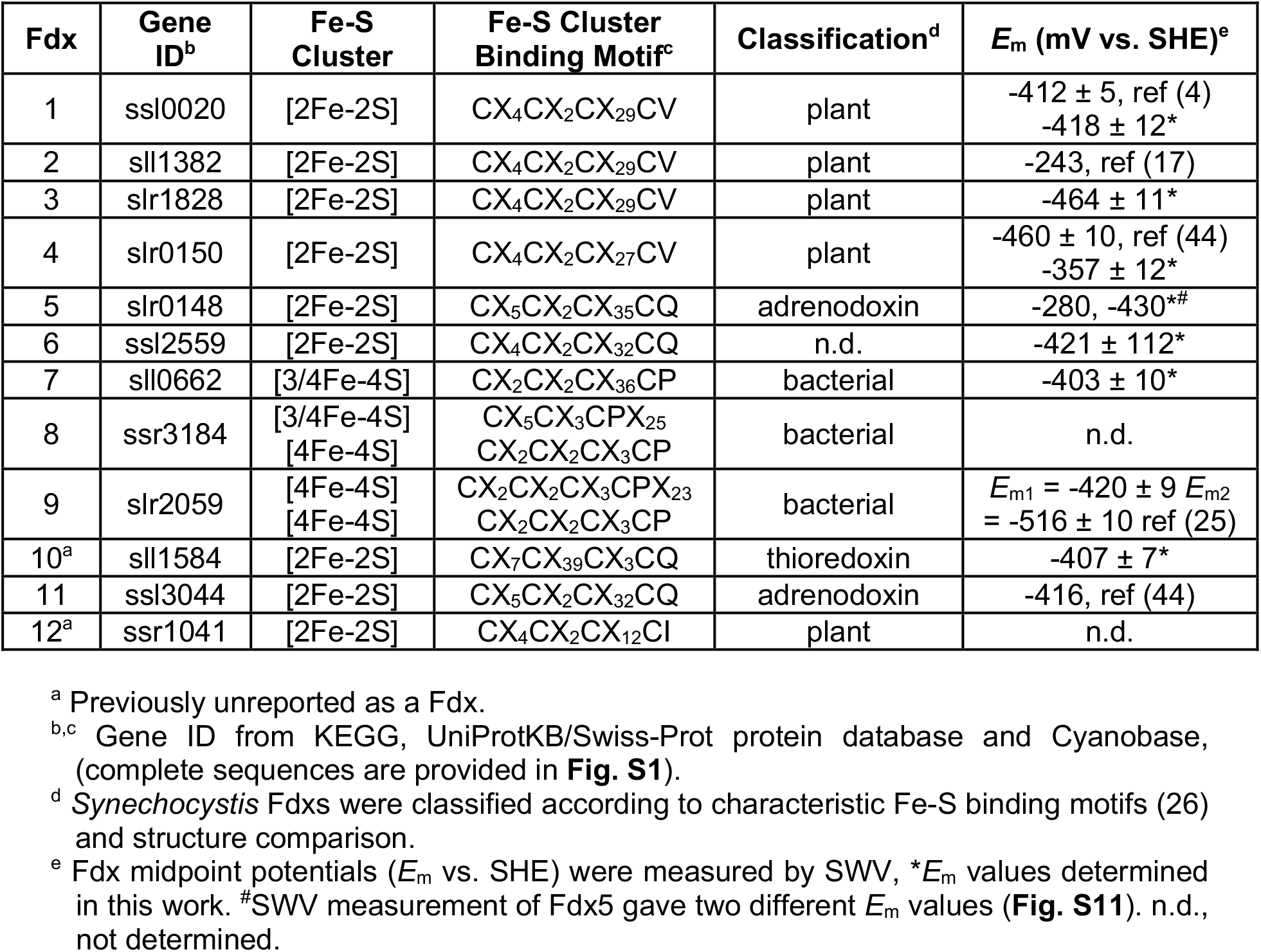
Data summary for the 12 Fdx and Fdx-like proteins from *Synechocystis*.

Fdx7, Fdx8 and Fdx9 form a distinct subgroup of bacterial Fdx-like proteins containing [4Fe-4S] or [3Fe-4S] clusters. Fdx7 is non-essential under standard growth conditions but becomes critical during oxidative stress, iron limitation, strong light, or exposure to selenate or cadmium (21-24). Previous studies suggested that Fdx7 and Fdx9 function in a coordinated stress response with the thioredoxin system (5, 22). Fdx8 appears to be essential for growth but has no assigned function (5, 20, 23). Based on the protein sequence, it was proposed to bind [3Fe–4S] and [4Fe–4S] clusters. Fdx9 is structurally unique based on a two-domain architecture and harboring two spin-coupled [4Fe-4S] clusters (*E*_m_ = –420 and –516 mV) (25). Fdx9 expression is induced under cold, high-light, or oxidative stress conditions (20), and deletion impairs photomixotrophic growth and metal-stress tolerance (5). Fdx9 supports photomixotrophic growth possibly by coupling electron transfer to PFOR (18) and is reported to interact *in vitro* with Flv3 based on two-hybrid assay experiments (26).

In this work, we evaluated the structural classifications and properties of the nine known, putative Fdxs and Fdx-like proteins, as well as three newly identified: Fdx10, Fdx11, and Fdx12. We used electron paramagnetic resonance spectroscopy and electrochemistry to refine the assignments of Fe-S cluster type and *E*_m_ values, respectively. We examined their expression patterns under a variety of growth conditions and tested reactivity with a few key physiological redox partners, Photosystem I (PSI), pyruvate ferredoxin oxidoreductase (PFOR) and nitrite reductase (Nir), offering new insights into how these proteins contribute to electron circuity and metabolic regulation (3, 5, 12, 15, 27). We integrate our findings with previously reported analyses to further resolve the roles of Fdx and Fdx-like proteins in the metabolism and biochemical function of electron transfer. Overall, the results advance a deeper understanding of how diversity in the *Synechocystis* Fdx proteins has evolved to support electron flow and maintain cellular redox homeostasis during adaptation to fluctuating environmental conditions.

## Materials and Methods

### Sequence retrieval

Previous studies reported on nine Ferredoxins in *Synechocystis* (5). Here we identified proteins Fdx10-12 in database searches on webpages (CYANOBASE, http://genome.kazusa.or.jp/cyanobase; CYORF, http://cyano.genome.jp/) (**Table 1**). Amino acid sequences were obtained from KEGG and UniProtKB/Swiss-Prot protein databases (**Fig. S1**).

### Computational structure analysis

Fdx structures and Fe-S cluster binding sites were predicted using AlphaFold2 (AF2) (28) and AlphaFill (29). The predicted structures were further refined using the AMBER force field. To assess structural and sequence similarities, proteins with significant homology to each ferredoxin were identified based on the Global Model Quality Estimate (GMQE) score from SWISS-MODEL, with scores above 0.5 considered significant (30). Protein surface charges were calculated using PyMOL and the APBS surface charge calculation plugin.

### Construction of expression and mutagenesis plasmids

*Synechocystis* ferredoxins were heterologously expressed in *E. coli*. The proteins were expressed from plasmids that added a TEV cleavage site containing linker (31) and either a tandem GST-6xHis-tag (Fdx1, Fdx2, Fdx3, Fdx4, Fdx5, Fdx6, Fdx7, Fdx9, Fdx10, Fdx11, Fdx12, IsiB) (32, 33) or a tandem MBP-6xHis-tag (Fdx6, Fdx8, Fdx9, Fdx10) to the C-terminus. To generate the GST-6xHis expression plasmids, the genes of interest were amplified from *Synechocytis* WT genomic DNA using the primers 1 to 26 (**Table S4**) and cloned into a previously described plasmid (33) that had been opened with *Nde*I and *Bam*HI. In some cases the obtained PCR fragments were subcloned into the pGEM®-T easy vector (Promega), while in others the fragments were directly inserted into the expression vector via Gibson cloning (34). To be able to generate the MBP-6xHis expression plasmids, some prior modifications to the GST-6xHis expression vector were required. By opening the vector with *Nde*I and *Eco*RI, only the pRSETA backbone was used. A fragment containing overlapping ends that allowed an initial insertion into the pBluescript SK+ plasmid (digested with *Kpn*I and *Sac*I) via Gibson cloning was ordered (Genscript, **Fig. S16**). The ordered fragment also contained the sequence encoding *Synechocystis* Fdx9, a 17-amino-acid (aa) linker (adapted from (31)) with an internal TEV cleavage site followed by a *Sma*I site, a 6xHis-tag, a stop codon (TAA), an *Eco*RV site, another stop codon (TAG) and an *Eco*RI site. It was excised again from the pBluescript vector using *Nde*I and *Eco*RI and inserted into the previously opened GST-6xHis expression vector yielding plasmid pMB0122. The MBP encoding sequence was amplified using primers 27 and 28 (**Table S4**) from the pRK793 plasmid (Addgene) and subcloned into the pGEM®-T easy vector (Promega). After excision with *Nhe*I and *Sma*I from the resulting pGEM®-T easy vector, the MBP-tag encoding sequence was inserted into the *Nhe*I and *Sma*I opened pMB0122 plasmid. For these expression vectors codon-optimized versions of previously difficult to express ferredoxins (Fdx6, Fdx8, Fdx9, Fdx10; Fig. S17), were ordered (Genscript) and could be inserted into the pMB0122 plasmid using the *Nde*I and *Bam*HI restriction sites. All expression plasmids were sequenced (Azenta). The construction of mutagenesis plasmids is described in **Supporting Text 1**.

### Protein expression and preparation

Table S2 lists the expression plasmids and strains that were utilized for each of the purified proteins. For Fdx6, Fdx9 and Fdx10 a mixture of cell cultures was used. The KRX strain (Promega) does not harbor an antibiotic resistance and protein expression was induced by the addition of 0.05% (w/v) L-rhamnose. The Rosetta™ 2 pLysS (DE3) strain (Novagen) harboring the pRARE2 plasmid is resistant to chloramphenicol (25 µg/ml) and expression was induced by the addition of 0.5 mM IPTG. The C41 strain harboring the pCodonPlus and pRKISC plasmids (35, 36) is resistant to chloramphenicol (25 µg/ml) and tetracycline (10 µg/ml) and expression was induced by the addition of 0.5 mM IPTG. The heterologous expression of Fdx8 was carried out in ΔiscR BL21 (37) to stimulate maturation of FeS-clusters in recombinant protein in a form that is biologically relevant. Fdx8 was expressed and purified under either aerobic or strictly anaerobic conditions. Fdx9 was expressed as described previously (25).

The expression plasmids used in this study conferred ampicillin resistance (100 µg/ml). A previously established method (32, 33) was modified and employed for protein preparation. In brief, a 5-ml LB starter culture was grown overnight at 37°C and diluted 1:100 in one or several 100-ml LB subculture the following morning. After approximately 3 h of growth at 37°C, 20 ml of the culture were used to inoculate 2L-flasks containing 1 L of LB. All cultures contained appropriate antibiotics and were grown aerobically. At an OD_600_ of around 0.7, protein production was induced by adding the appropriate inducing agent and 1mM ferric ammonium citrate prior to a shift to 20°C. Cells were harvested the following morning (15 min, 4°C, 5.000 x g) and resuspended in lysis buffer (50 mM NaPO4 pH=7.0, 250 mM NaCl) for breakage by sonication (Sontrode MS73 (Bandelin); 8 repeats of 20 s on (70% cycle, 70% power) and 20 s off). The supernatant obtained after ultracentrifugation (45 min, 4°C, 100.000 x g) was incubated for 1 h at 4°C with TALON® Superflow™ resin (Takara). After the incubation period, the resin was washed with 20 column volumes (CV) of lysis buffer or until the wash solution became clear and colorless. Protein elution was performed with 5 CV of elution buffer (50 mM NaPO4 pH=7.0, 250 mM NaCl, 250 mM imidazole). The eluted proteins were dialyzed overnight in 50 mM Tris pH=8.0, 100 mM NaCl in the presence of TEV-His protease (His-tag purified from pRK193 (Addgene) (38). The following day, the protein was incubated again with TALON® Superflow™ resin (Takara) and the flow-through was collected, concentrated and loaded onto a HiLoad™ 26/60 Superdex™ 75 prep grade (Cytiva) using a previously established method (33). The run was monitored at 280 nm, 2-ml fractions were collected and the final protein concentrations were determined using ROTI®Nanoquant assay (Roth) according to the manufacturer’s instructions.

### Protein analyses

To assess the purity of heterologously expressed protein after purification, some sample was mixed with 2x Laemmli sample buffer, boiled for 5 min at 95°C and centrifuged in a microfuge (5 min, max speed). Roughly 5 µg of the purified protein were then loaded per lane and separated on a 15% (w/v) polyacrylamide BisTris gel using MES running buffer (250 mM MES, 250 mM Tris, 5 mM EDTA, 0.5 % (w/v) SDS). To probe ferredoxin expression levels, aliquots corresponding to 50 ml of the respective cell cultures were resuspended in 500 µl of ACA buffer (750 mM ε-aminocaproic acid, 50 mM BisTris/HCl, pH 7.0, 0.5 mM EDTA). Cells were broken by vortexing with ∼200 µl glass beads (0.17–0.18 µm diameter; Sigma) for 2 min at full speed at 4°C. Glass beads and cell debris were then pelleted by centrifuging in a microfuge at full speed for 1 min. The whole cell extract (WCE) was centrifuged again at 20,000g for 20 min, the supernatant was kept as the soluble extract and its protein concentration was determined using the ROTI-Quant protein Bradford assay according to the manufacturer’s instructions (Carl Roth). Due to the fact that cell breakage was different for different samples and to ensure equal loading within one dataset (growth condition), varying amounts of proteins (autotroph (10 µg), mixotroph (10 µg), heterotroph (7.5 µg), mixotroph with arginine as nitrogen source (10 µg), nitrate depletion (1 µg) and Fe depletion (10 µg)) were loaded per lane on a 17.5% (w/v) polyacrylamide BisTris gel using MES running buffer (250 mM MES, 250 mM Tris, 5 mM EDTA, 0.5 % (w/v) SDS). Gels were Coomassie-stained or electroblotted onto nitrocellulose membrane. For immunoblotting, 5% (w/v) milk powder in 1x PBS-T was used as blocking solution and all washing steps were performed with 1x PBS-T. Immunoblot analyses were performed using THE™ His-tag antibody (Genscript) and horseradish peroxidase-conjugated secondary antibodies (anti-mouse, Cytiva).

### UV/Visible absorption spectra

The spectra of the different ferredoxins and flavodoxin were measured in the range from 300 to 600 nm using a Cary-60 UV-Vis photometer (Agilent, CA, USA). In a total volume of 1 ml of 100 mM Tris pH=8 a protein concentration that gave a reading in its maximum of at least 0.3 was used. After subtraction of the baseline this was used as the oxidized spectrum. After addition of 40 µl of freshly prepared 100 mM sodium dithionite to the same sample the reduced spectrum was recorded. Since dithionite gives a strong reading below 380 nm only the spectra above this wavelength are shown.

### Electron paramagnetic resonance experiments

EPR samples of reduced ferredoxins 1-4, 7, and 10-11 (100 µM final concentration) were prepared in 25 mm NaPO_4_ pH 7.0, 50 mM NaCl, 5% glycerol under anaerobic conditions in an MBRAUN glovebox (nitrogen atmosphere). Samples were reduced with NaDT (5 mM final concentration) and loaded into EPR tubes (Wilmad 707-SQ-250m), which were sealed with septa and frozen in liquid nitrogen. Ferredoxin 8 was reconstituted according to the procedure previously described for Fdx9 (25). A reduced sample for EPR was prepared by reduction with 10 mM NaDT in 50 mM Tris pH 8.3, 200 mM NaCl, 5% glycerol. EPR was collected on a Bruker Elexsys E500 spectrometer as previously described (25). Spectra were processed in OriginPro 2024b by baseline correction through subtraction of a spline function and background subtraction of buffer. Simulations of the EPR spectra were carried out in the EasySpin toolbox (39) using the ‘esfit’ least-squares fitting algorithm and EasySpin ‘pepper’ function in MATLAB R2023b.

### Expression level analysis

*Synechocystis* cultures to probe ferredoxin expression levels were grown under the following conditions: autotroph (BG11; start OD_750_ = 0.3; 5 days), mixotroph (BG11, 10 mM glucose; start OD_750_ = 0.2; 2 days), heterotroph (dark; BG11, 10 mM glucose; start OD_750_ = 0.2; 2 days), mixotroph with arginine as nitrogen source (BG11_0_, 10 mM glucose, 5 mM arginine; start OD_750_ = 0.15; 3 days), nitrate depletion (BG11_0_; start OD_750_ = 0.15; 3 days) and Fe depletion (BG11 -Fe; start OD_750_ = 0.2; 3 days) (40). In each case, 50-ml pre-cultures with appropriate antibiotics were started four days ahead of the experiment and cultivation started after washing the pre-cultures with the target growth medium (without antibiotics) and adjusting the start OD_750_ to the indicated values. Cell growth occurred in one Kniese tubes per strain (WT, Fdx1-6xHis to Fdx11-6xHis) in a volume of 150-200 ml liquid culture with constant ambient air bubbling at 28°C and 50 μE m^-2^ s^-1^ light. Cells were harvested after the indicated durations and subjected to protein analyses.

### Square wave voltammetry

Protein-film square wave voltammetry (SWV) was performed in an MBraun anaerobic chamber according to previously published protocols (41). Square wave voltammetry (SWV) measurements were performed in a standard three-electrode cell using a pyrolytic graphite edge (PGE) working electrode, an Ag/AgCl (saturated KCl) reference electrode, and a platinum wire counter electrode. The working electrode was polished before each experiment, and background signals were recorded in buffer without protein. Protein films were incubated on the electrode for at least 10 minutes prior to measurements in a buffer containing 150 mM HEPES, 200 mM NaCl, and 5% glycerol at pH 8. The same buffer was used as the electrolyte. SWV was conducted using a CH Instruments 630C potentiostat with a 0.001 V step size, 0.025 V amplitude, 10 Hz frequency, and 1 μA/V sensitivity. Current and potential were simultaneously sampled at 1 MHz. All potentials were converted to values versus the standard hydrogen electrode (SHE) by adding 199 mV to the Ag/AgCl reference.

### NanoLuc interaction studies

Strains to assess the interaction between nitrite reductase and Fdxs were grown under autotrophic growth conditions. Cells were pelleted by centrifugation and the OD_750_ was normalized to 1. 200 µl of normalized cell culture were transferred into three wells of a white 96-well plate with a transparent bottom and 50 µl of 25 µM Coelenterazine as substrate were added. After transferring the plate into an Infinite M Plex plate reader (Tecan) and reaching 30°C in the inner chamber, OD_750_ was measured followed by a recording of the luminescence for 30 min (integration time 2s). The luminescence signals were averaged per well and normalized according to measured OD_750_. Then the signal was average for each strain and plotted.

### P700^+^ Reduction Kinetics

Electron transfer from PSI to various Fdxs was assessed by monitoring P700^+^ re-reduction kinetics using a Dual-KLAS/NIR instrument (42). Assays (1 ml) contained 500 nM PSI (based on Chl a content), 500–2,500 nM Fdx, and an oxygen-scavenging system (10 mM glucose, 20 U glucose oxidase and 25 U catalase in activity buffer (25 mM Tris-HCl buffer pH 8.0, 10 mM CaCl_2_, 10 mM MgCl_2_ and 0.03 % (w/v) beta-dodecyl-D-maltoside). Following a 5-minute dark incubation, the sample was subjected to a 10 ms multiple-turnover light pulse (13,600 µE m^-2^s^-1^). In the absence of an acceptor, P700^+^ re-reduction is rapid due to charge recombination, i.e. successful electron transfer to an Fdx slows this re-reduction rate. PSI alone (no transfer) and PSI with 1 mM methyl viologen (complete transfer) served as benchmarks. The P700^+^ re-reduction slope was calculated over the first 100 ms post-illumination. Slopes were normalized against signal amplitude (pre-vs. post-pulse) and compared to the PSI-only control to determine the percentage of electron transfer.

### PFOR activity assay

To assay potential interactions between the various ferredoxins and the pyruvate ferredoxin oxido reductase (PFOR), the method reported previously (18) was adapted. Briefly, an assay with a final volume of 1 ml containing 4.5 mM ferredoxin, 270 μM PFOR, 0.5 mM coenzyme A, 10 mM pyruvate, 5 mM thiamine pyruvate phosphate as well as 40 mM glucose, 40 U glucose oxidase and 50 U catalase (to maintain anaerobiosis) in 100 mM Tris-HCl buffer pH 8.0 was incubated for 30 min at room temperature (25°C) while absorption at 460 nm was monitored using a Cary 60 UV-Vis photometer (Agilent). The assay was started by the addition of The assay was started by the addition of the respective ferredoxin at concentrations up to 100 µM.

## Results

### Sequence and structural comparisons of ferredoxins Fdx1-Fdx12

Fdx1 to Fdx9 were previously described and classified (5). Three additional Fdxs, Fdx10 (sll1584) (15), Fdx11 (ssl3044) (12), and Fdx12 (ssr1041) (this study) were identified (Table 1). Fdxs typically coordinate either [2Fe-2S] or [4Fe-4S] cluster(s) (3, 27, 43). They are further classified based on amino acid sequence, protein structure, and the cysteine-rich motifs that coordinate the Fe-S cluster(s) (**Table 1, Figs. 1** and **S1**) (3, 27).

**Figure 1.**
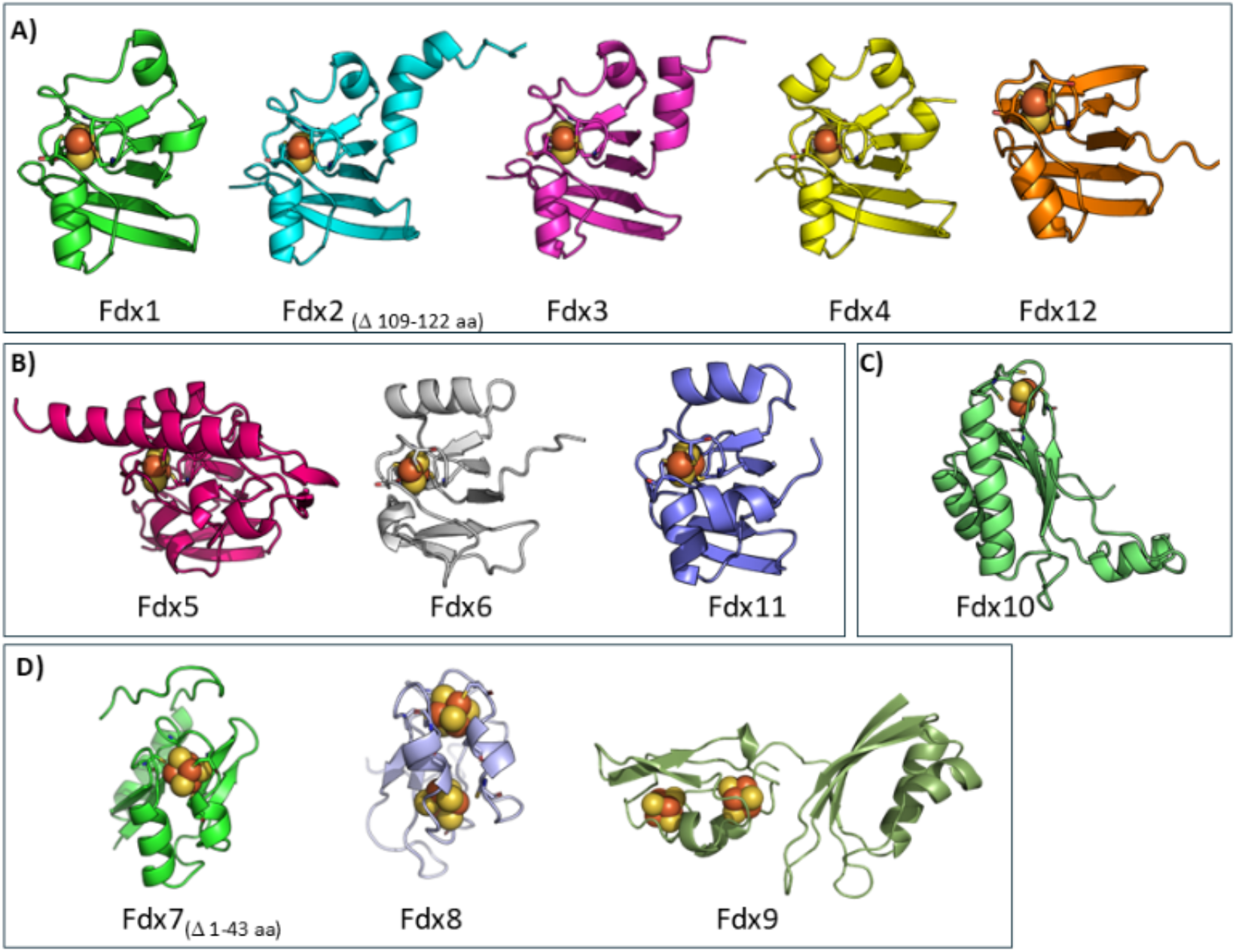
Comparison of *Synechocystis* Fdx protein structures and classifications. The comparisons reveal the structural variability of *Synechocystis* Fdxs. **A)** [2Fe-2S] plant-type, **B)** [2Fe-2S] cluster adrenodoxin-type, Fdx5 and Fdx11; Fdx6, undetermined. **C)** [2Fe-2S] thioredoxin-like. **D)** [4Fe-4S] bacterial-type. For Fdx1 an experimental structure is available (PDB ID: 1OFF). Structures for [2Fe-2S] cluster Fdxs were predicted using AlphaFold2. The clusters were placed using AlphaFill and minimized using the AMBER forcefield. The Fe-S clusters are shown as spheres while the protein structures are shown as a cartoon representation. Placement of [4Fe-4S] clusters in Fdx7 and Fdx8 were determined based on Pymol generated models using PDBs of structural homologues; PDB ID: 1FXR and PDB ID: 1H98. Truncated versions are shown for Fdx2 (Δ109-122) and Fdx7 (Δ1-43), because the respective segments were predicted with low confidence by AlphaFold. Fdx5 was modeled using the short sequence (**Fig. S1**). The Fdx9 model structure was previously reported (25).

As Fdx1 is the only *Synechocystis* Fdx with experimentally resolved structure (45), protein structures of the other 11 Fdxs were obtained *in silico* (**Fig. 1**). Similarities between Fdx1 and the other plant-type *Synechocystis* Fdxs were quantified using Root Mean Square Deviation (RMSD) calculations (46). Calculated RMSD values for Fdx2, Fdx3, Fdx4 and Fdx12 were 0.918, 0.495, 0.457, and 2.806 Å respectively, with values below 2 suggesting significant structural homology. Plant-type [2Fe-2S] Fdxs typically exhibit high sequence and structural homology (3), as holds true for Fdx1, Fdx2, Fdx3, Fdx4, and Fdx12 (**Fig. 1A**). They adopt a highly conserved β-grasp fold, comprising a mixed β-sheet with five β-strands and one to three adjacent α-helices (3). The [2Fe-2S] cluster is coordinated by four conserved cysteines in a CX_4_CX_2_CX_n_C motif and further stabilized by a network of hydrogen bonds.

Structures and sequences of Fdx5 and Fdx11, resemble adrenodoxin-type [2Fe-2S] cluster Fdxs (**Figs. 1B** and **S2**) which are versatile proteins that function as soluble electron carriers in a plethora of reactions. They typically consist of a conserved core domain essential for electron transfer and a variable domain usually including a hydrophobic loop and a flexible C-terminus for the recognition and interaction with their protein partners (47). Fdx5 is the largest protein in the group exhibiting the most complex structure, including a flexible domain with a 20 amino acid long α-helix, as well as a less accessible cluster (**Figs. 1B** and **S2**). Fdx6 seems to have an atypical [2Fe-2S] cluster binding motif, with structural characteristics resembling both plant and adrenodoxin type. EPR characterization of this Fe-S cluster is discussed later but shows features more in line with a plant-type [2Fe-2S] cluster.

Fdx10 is a thioredoxin-like Fdx with a structure that is distinct from other Fdxs and features a highly conserved thioredoxin fold, consisting of a central four-stranded β-sheet surrounded by three α-helices (**Fig. 1C**) (48). Notably, Fdx10 contains an atypical Fe-S binding sequence, C_32_C_33_X_6_C_40_C_41_X_38_C_80_X_3_C_84_Q, where one cysteine from each Cys-Cys pair, Cys32 and Cys40, coordinates the [2Fe-2S] cluster. The structure suggests that the adjacent cysteines, Cys33 and Cys40, may play a role in stabilizing the cluster or modulating its reactivity, an intriguing possibility that warrants further investigation. The remaining cysteines (Cys17 and Cys64) are far away from the [2Fe-2S] cluster, in positions that could allow for the formation of S-S bonds and a thioredoxin-like function, and may be involved in redox sensing and regulation (49). The function of adrenodoxin and thioredoxin-type Fdxs remain poorly understood in cyanobacteria, and our analysis provides a first step toward filling this gap by revealing their distinct structural features and potential roles in redox sensing and protein regulation.

The bacterial-type [3Fe–4S]/[4Fe–4S] group includes Fdx7, Fdx8, and Fdx9, each exhibiting βαββαβ secondary structure motifs typical of this family. Fdx7, among the largest *Synechocystis* putative Fdxs, possesses extended N- and C-terminal segments with intrinsic flexibility, possibly mediating complex formation with partner proteins (**Table 1, Figs. 1** and **S3**). The position of the cysteines confirms binding of a single [4Fe-4S] cluster, while an additional cysteine contributes Fe-S cluster stabilization through hydrogen bonds. Based primarily on its amino acid sequence and predicted structural features, Fdx8 is most consistent with the assignment as a [3Fe-4S], [4Fe-4S] di-cluster Fdx (14). Notably, the canonical ligand for a [4Fe-4S] cluster is replaced by a non-polar valine residue, indicating the absence of a fourth, conventional iron-coordinating cysteine. This is supported by sequence alignments with representative 2x[4Fe-4S] or [3Fe-4S], [4Fe-4S] di-cluster Fdxs (**Fig. S4**). A particularly intriguing observation regarding the cluster composition of Fdx8, discussed in more detail later in the text, was an apparent discrepancy between the sequence-based prediction of a [3Fe-4S] and a [4Fe-4S] cluster and the experimentally observed presence of 2x[4Fe-4S] clusters. Structural modeling of Fdx8 (**Fig. S5**) suggests that the amide nitrogen of the valine residue is positioned near the [3Fe-4S] cubane, in a location that could potentially serve in coordinating a fourth iron atom to stabilize a [4Fe-4S] cluster (**Fig. S5**). However, this hypothesis requires further experimental verification (5). Fdx9 features an unusual two-domain structure, with one domain resembling a clostridial-type 2x[4Fe-4S] Fdx, while the second domain resembles a Nil-like structure with unidentified function (25).

### In silico analysis of Fdx1-12 surface charges

The spatial arrangement of charged amino acids on protein surfaces is known to affect the electrostatic contribution to the formation of protein-protein interactions, which are necessary for electron transfer to a partner enzyme. The strength of the electrostatic contribution directly correlates to the number of complementary surface charges between a protein-protein pair. To assess the distribution of charged residues on the surfaces of Fdx1-Fdx12, we performed an *in silico* analysis on each of the structural models (**Fig. 2**).

**Figure 2.**
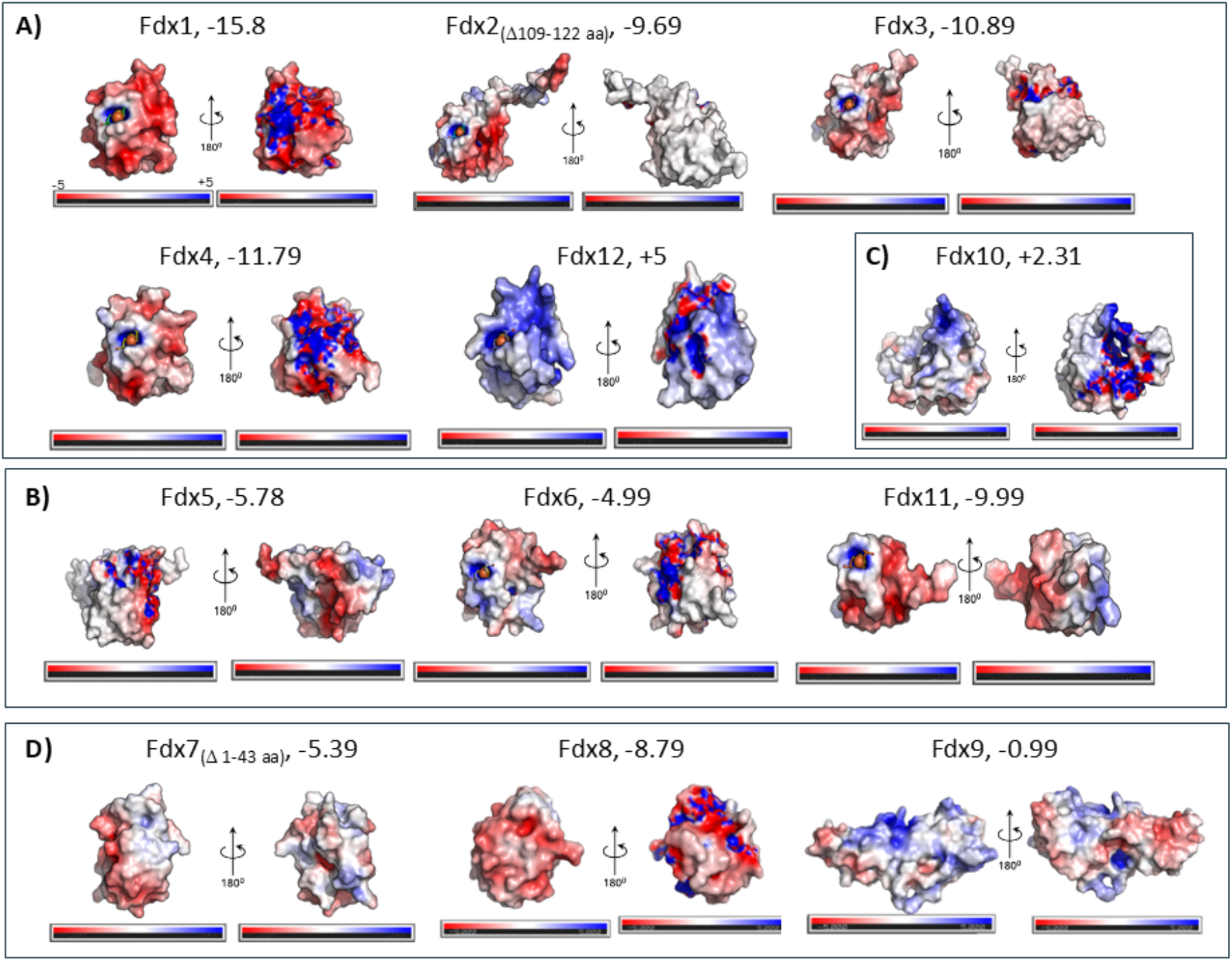
A comparison of the surface charges of Fdxs 1-12 calculated for pH 7: A) [2Fe-2S] plant-type, B) [2Fe-2S] adrenodoxin-type and Fdx6, C) [2Fe-2S] thioredoxin-like type, D) [4Fe-4S] bacteria-type. Surface charges for Fdx9 were reported previously (25). The surface charge predictions were obtained in Pymol using the APBS surface charge calculation plugin. The blue regions indicate positively charged residues, the red regions indicate negatively charged residues, white regions indicate neutral surface. Due to the regions with poorly defined AlphaFold structure for Fdx2 and Fdx7, calculations may misrepresent the overall surface charges. Fdx5 was modeled using the short sequence (**Fig. S1**).

Plant-type Fdxs generally exhibit acidic surfaces that favor interaction with positively charged partners involved in photosynthetic electron transfer and downstream metabolism (35). Consistent with this, Fdx1 and Fdx4 show highly acidic surfaces, whereas Fdx2 and Fdx3 combine acidic and hydrophobic regions, implying alternative binding partners. Fdx5, and Fdx6 display mixed acidic and basic patches typical for adrenodoxin-type Fdxs (36), supporting versatile electrostatic and hydrophobic interactions. Among the bacterial-type Fdxs, Fdx7 and Fdx8 are overall acidic. In contrast, Fdx10 and Fdx12 show basic surfaces, unusual for proteins with Fdx function (6, 25, 50).

### Comparative biophysical characterization of Fdx1-Fdx11

*Synechocystis* Fdx1-Fdx11 and IsiB (flavodoxin) were over-expressed in *E. coli* and purified (**Table S2, Fig. S6**). To assess the purity and integrity of these proteins and to further characterize their Fe-S clusters, UV-Vis absorption spectra (**Figs. S7** and **S8)** and EPR spectra (**Figs. 3** and **S9** and **Table S3**) were recorded.

**Figure 3.**
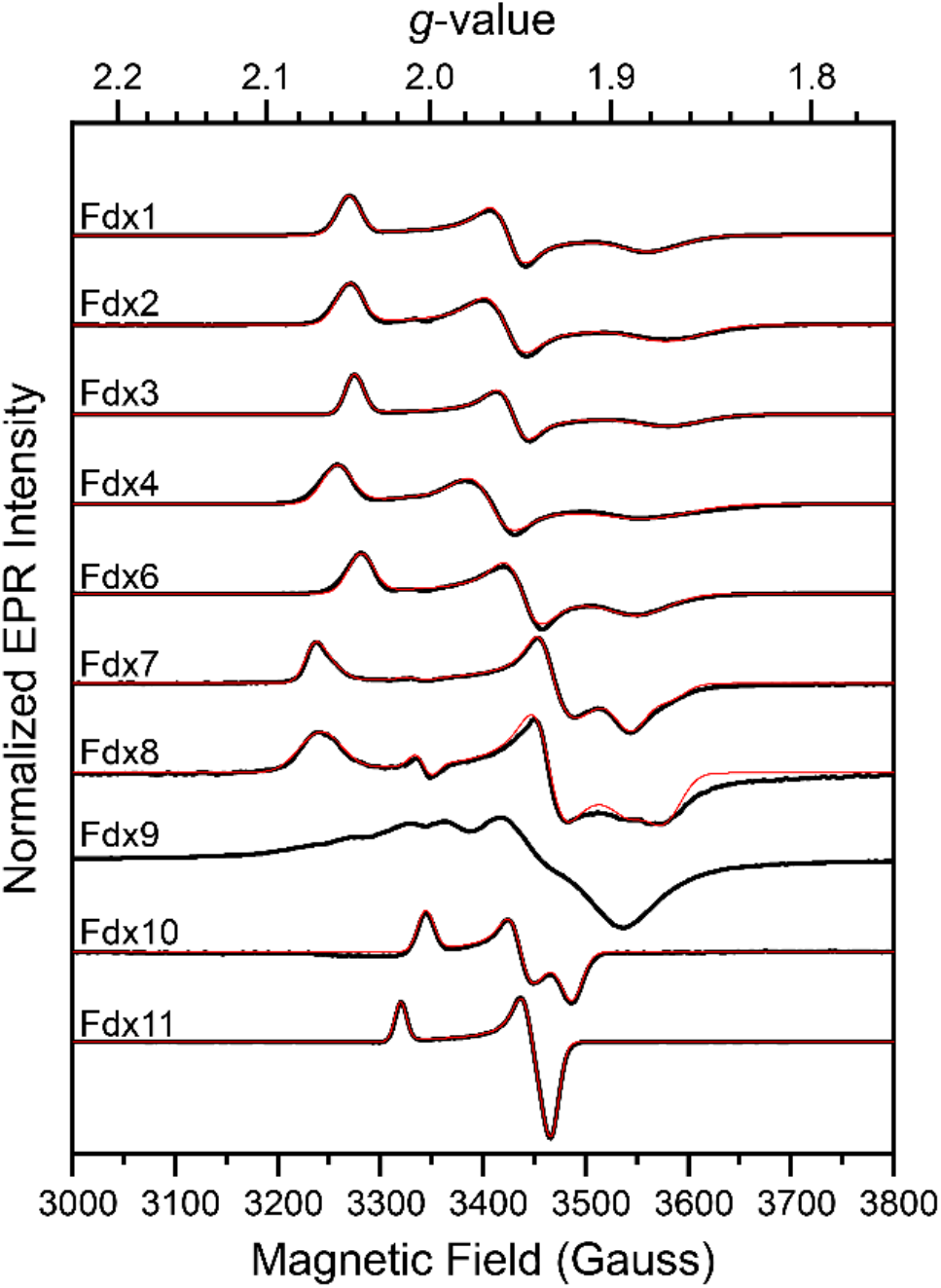
Electron paramagnetic resonance (EPR) spectra of reduced Fdx1-11. Fdx samples were reduced with sodium dithionite. Experimental traces are shown in black and simulated traces are shown in red. For Fdx5 no EPR signals could be observed. Fdxs 8 and 9 were reconstituted with iron and sulfide under anaerobic conditions. A simulation for Fdx9 was omitted due to spectral complexity from spin-spin coupling (25). All spectra were recorded at 1 mW microwave power. Fdx1 to Fdx4 and Fdx6 to Fdx8 were recorded at 20 K, Fdx9 at 10 K, Fdx10 and Fdx11 at 30 K.

A comparative analysis of UV-Vis spectra for Fdx1, Fdx7, and Fdx8 with oxidized [2Fe−2S] and [4Fe−4S] clusters, respectively, illustrates characteristic spectral features of different cluster types (**Fig. S8A**). All *Synechocystis* [2Fe-2S] cluster Fdxs exhibit similar UV-Vis absorption profiles, with characteristic peaks between 414–426 nm and 451–470 nm (**Fig. S7**). The primary distinction between spectra from different [2Fe-2S] cluster Fdxs lies in the relative intensities of the peaks at 420 nm and 460 nm. Fdx7 and Fdx8 spectra display broader, less resolved peaks around 420 nm, a feature characteristic of oxidized [4Fe-4S] clusters. The UV-Vis spectra of all reduced Fdxs show lower intensity and broader peaks, consistent with reduced Fe-S clusters. Remarkably, Fdx5 could not be reconstituted and reduced under the conditions that were applied to other Fdxs (**Fig. S7**), possibly due to inaccessibility of the [2Fe-2S] cluster to reducing reagent.

Notably, anaerobically purified Fdx8 exhibits spectral features that are characteristic of [4Fe-4S] clusters (**Fig. S8**) (51) despite its sequence suggesting a more likely [3Fe-4S] [4Fe-4S] configuration (**Figs. S4** and **S5**). Upon exposure to air, broad peaks appear around 315 nm and 410 nm, indicative of the formation of oxidized [3Fe-4S] clusters (51). These experimental results suggest that one of the Fdx8 clusters, upon exposure to oxygen, transforms from a [4Fe-4S] to a [3Fe-4S] cluster (**Fig. S8**) (25). The physiological significance of an interchangeable [3/4Fe-4S] cluster and possible oxygen sensing function remains to be investigated.

### Electron paramagnetic spectroscopy

A survey of the EPR spectral properties of reduced Fdxs from *Synechocystis* was carried out to aid in the assignment of the Fe-S cluster type, and to inform on the various cluster geometric and magnetic properties (**Figs. 3** and **S9, Table S3**).

Reduced Fdxs1-4 exhibit similar spectra represented by rhombic *S* = 1/2 signals with calculated rhombicity values typical for plant-type [2Fe-2S] cluster Fdxs (**Table S3**) (4, 52, 53). Slight variation in the degree of rhombicity is indicative of small differences in the distortion of the Fe^2+^ environment for the mixed-valent (Fe^2+^-Fe^3+^) [2Fe-2S] cluster (54). Likewise, reduced Fdx6 displayed a rhombic *S* = 1/2 signal reflective of a [2Fe-2S] cluster, however it is difficult to distinguish whether the signal reflects either a plant or adrenodoxin-type Fdx. The signal lacked axial-type symmetry that is a signature of other adrenodoxin-type Fdxs (55, 56), however it displayed a slightly lower degree of rhombicity (75%) compared to plant-type Fdxs 1-4 (**Fig. 3, Table S3**). Reduced Fdx10 and Fdx11 display S = 1/2 signals with *g*-value anisotropy typical for thioredoxin-like (57) and adrenodoxin-type [2Fe-2S] cluster Fdxs, respectively (55, 56). We have not observed an EPR signal for reduced Fdx5, even after reconstitution, which is consistent with UV-Vis experiments (**Table S3**).

Reduced Fdx7 and Fdx9 (shown here and described in (25), respectively) display *S* = 1/2 signals typical for bacterial-type Fdxs that contain [4Fe-4S] clusters (58, 59) (**Fig. 3**). However, the spectrum of Fdx7 is more complex than expected for a single, reduced, [4Fe-4S] cluster as has been previously assigned (5, 24) and spectral fits identified two overlapping signals with different rhombicity, *g*-values and relative contributions (**Table S3**). The functional relevance of this spectral complexity is not immediately clear but may be indicative of conformational variability in reduced Fdx7 leading to heterogeneity in the Fe-S cluster electronic structure. Compared to Fdx7, reduced Fdx9 displays an even more complex spectrum that is characteristic of 2x[4Fe-4S] magnetically interacting clusters. In this case, spin-spin coupling between the clusters is thought to give rise to the multitude of features in the *g* = 2 region, as well as line-shape broadening, as described previously (25).

Reduction of anaerobically prepared and reconstituted Fdx8, displayed a complex spectrum that could be simulated as two overlapping rhombic components consistent with two reduced [4Fe-4S] clusters (**Figs. 3** and **S9**). As noted above, the Fe-S cluster binding motifs of Fdx8 are predicted to coordinate a [3Fe-4S] cluster and a [4Fe-4S] cluster (5). Conversion of the [3Fe-4S] clusters into [4Fe-4S] clusters in the reconstituted protein may explain this difference. This type of cluster conversion has been reported in detail for other [3Fe-4S], [4Fe-4S] di-cluster Fdxs prepared under reducing conditions (60-65). Electronic interaction between the clusters may also contribute to the complexity of the spectrum, and for other di-cluster Fdxs, spin-spin coupling between clusters has been shown to lead to additional complexity of the *S* = ½ region as observed here for Fdx8 (66, 67). This phenomenon may necessitate more advanced simulation, and the possible spin-spin coupling could account for features in the *g* = 1.9 region, as well as broadening on the wings of the spectrum, that were not able to be replicated exactly in the simulation. Additional EPR experiments that investigated the effects of oxygen on Fdx8 suggest that one of the clusters, most likely coordinated by the three-cysteine motif, can alternate between a [3Fe-4S] and [4Fe-4S] (**Table S4, Fig. S9**). Experiments on Fdx8 specifically enriched in either [3Fe-4S], [4Fe-4S] or 2x [4Fe-4S] clusters are needed to determine the functional relevance of each type. For example, whether these cluster changes are involved in modulating the electron transfer reactivity of Fdx8 with redox partners.

### Comparative in vivo expression of Fdx1-Fdx11

To gain further information on the regulation of Fdx protein expression levels under different growth conditions we constructed individual strains that expressed His-tagged versions of each Fdx (Fdx1 to Fdx11). Equivalent culture samples were separated by electrophoresis and used for immunoblotting with an antibody to the 6xHis-tag and visualized in side-by-side comparison (**Fig. 4** and **S10**).

**Figure 4.**
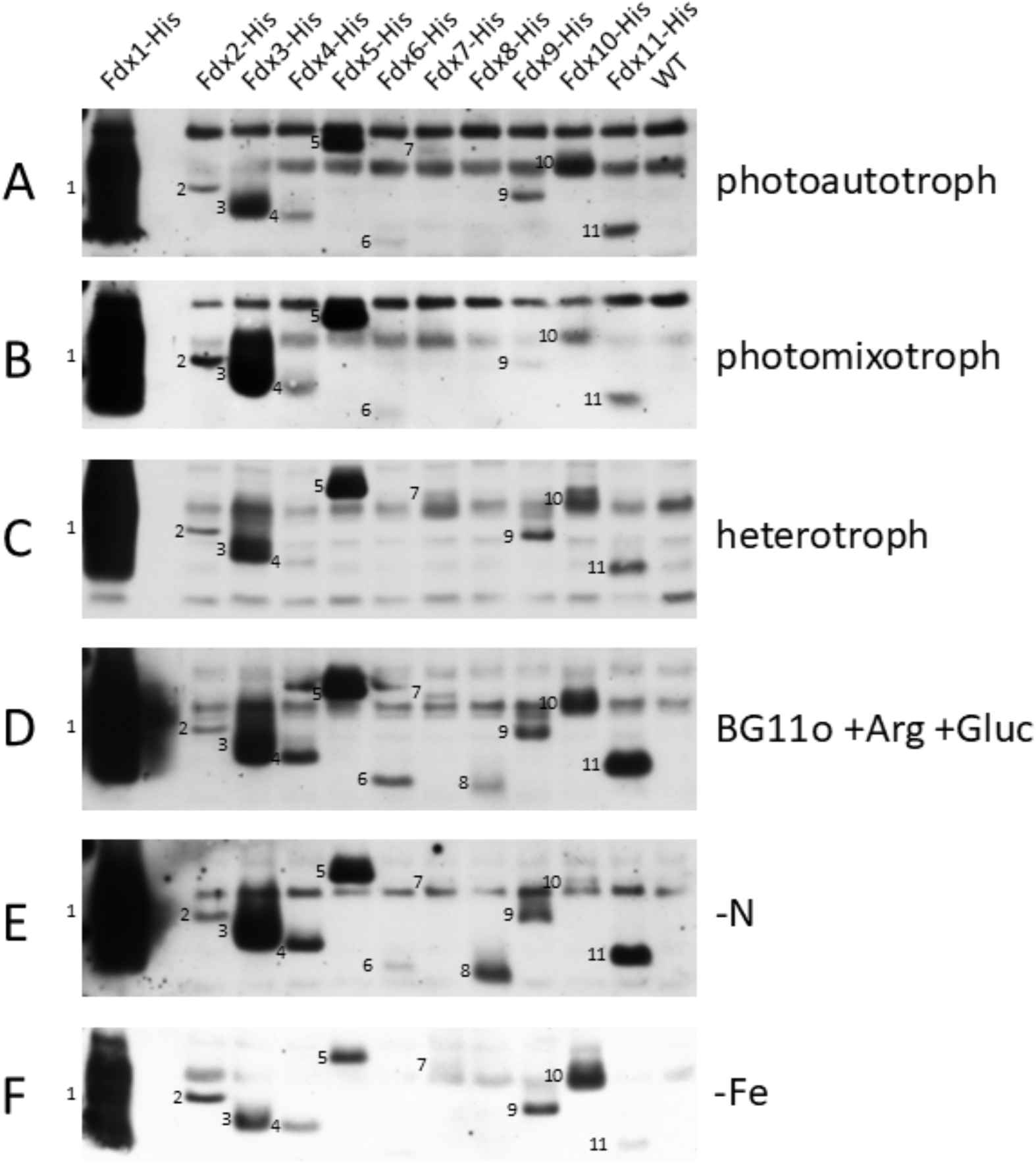
Expression levels of His-tagged Fdxs 1-11 under various growth conditions assessed by immunoblotting with an α-His antibody. Fdx reactive bands are marked with their respective number.

We tested photoautotrophic, photomixotrophic, and heterotrophic conditions as well as nitrate and iron limitation. In addition, growth under arginine and glucose (BG11_0_+Arg+Glc) culture conditions was tested as this is promotes a highly reducing metabolism that requires HoxEFUYH expression for growth (68). All Fdxs are detectable in at least one of the probed growth conditions. In all conditions Fdx1 is by far the most dominant isoform as reported previously (5) followed by Fdx3 and Fdx5. Whereas Fdx2, Fdx3, Fdx4, Fdx5, Fdx9, Fdx10 and Fdx11 are expressed under all conditions but at different levels, the Fdx6, Fdx7 and Fdx8 are expressed only at low-levels or under detection limits under these conditions. Fdx8 and Fdx4 expressions were highest in cells grown under nitrate depletion (-N) or grown photomixotrophically with arginine as nitrogen source (BG11_0_ +Arg +Gluc) (**Fig. 4**, panels **C** and **D**). Fdx4 and Fdx5 show distinctive expression levels even though they are organized in an operon and co-transcribed (69). The expression level of Fdx5 was at similar levels under all growth conditions.

Fdx10 and Fdx11 are expressed under photoautotrophic conditions and may be involved in stress adaptation, since Fdx10 and Fdx11 have been shown to be up-regulated under low temperature and Fdx11 is upregulated under high-light and Fe-limitation (19). Here we also observed that Fdx10 is up-regulated in cultures grown under heterotrophic, Fe-limitation, or in BG11_0_+Arg+Glc (**Fig. 4**, panels **C, F** and **D**). Fdx11 was also up-regulated in cells grown on BG11_0_+Arg+Glc or under N-limitation (**Fig. 4**, panels **D** and **E**).

### Determination of redox potentials

The formal mid-point redox potentials (*E*_m_) of a Fdx contributes to the donor-acceptor free-energy for electron transfer with a protein partner. The *E*_m_ values of Fdx1-Fdx11 were investigated using square wave voltammetry (SWV), with the purified proteins immobilized on a pyrolytic graphite edge (PGE) working electrode (**Table 1, Fig. S11**). For all Fdxs, except Fdx9, a single redox peak was observed (**Fig. S8**) (25).

The *E*_m_ values of Fdxs1-11 were found to span a wide range of potential from -243 mV to -520 mV vs SHE (**Table 1**). The SWV analysis of Fdx5 produced an individual reduction potential peak that shifted in potential with repeated SWV scans, which suggests it may have reorientated on the PGE surface, adopting different structures with different *E*_m_ values. The *E*_m_ value of Fdx4 differed by approximately 100 mV when measured on a PGE versus a gold electrode. As the more negative *E*_m_ value is closer to values reported for a Fdx4 homolog measured in solution (54) this value will be considered further. It is important to note that *E*_m_ values obtained via protein film voltammetry can vary depending on the electrode material and may differ from those measured in solution. Such shifts in *E*_m_ values are likely protein-specific and may reflect differences in surface binding and conformational states on the electrode surface (3, 70).

### In vivo investigation on the interaction of nitrite reductase NirA with Fdxs1-12

Nitrite reductase (NirA, slr0898) is a redox enzyme that functions in nitrogen metabolism and based on relative *E*_m_ values, may couple reactivity to electron transfer with one of the Fdxs (**Fig. S12**). We tested this by examining the ability of a Fdx to form a binding interaction with NirA using a NanoLuc luminescence complementation assay (Nanobit). The large subunit (nLucL) was fused to NirA and the small subunit, nLucS was fused to a Fdx, where a binding interaction results in a luminescence signal (71). When Fdx1-11, nLucS fusions were expressed from their native promoters in strains grown under photoautotrophic conditions, only Fdx1-nLucS produced a luminescence signal indicating an *in vivo* interaction with NirA-nLucL (**Fig. S13**). Since Fdx expression levels are highly regulated by growth conditions (**Fig. 4**), we constructed strains to bypass native regulatory control using rhamnose-inducible expression of Fdx-nLucS as well as IsiB-nLucS, (see Methods). Under the same growth conditions, the results also showed interactions between NirA-nLucL and Fdx4-nLucS and Fdx5-nLucS (**Fig. 5**). These binding interaction results with NirA and demonstration of Fdx4 and Fdx5 expression being elevated under nitrogen-limiting conditions or nitrogen source (**Fig. 4**) indicate a function in nitrogen metabolism by mediating electron transfer with NirA.

**Figure 5.**
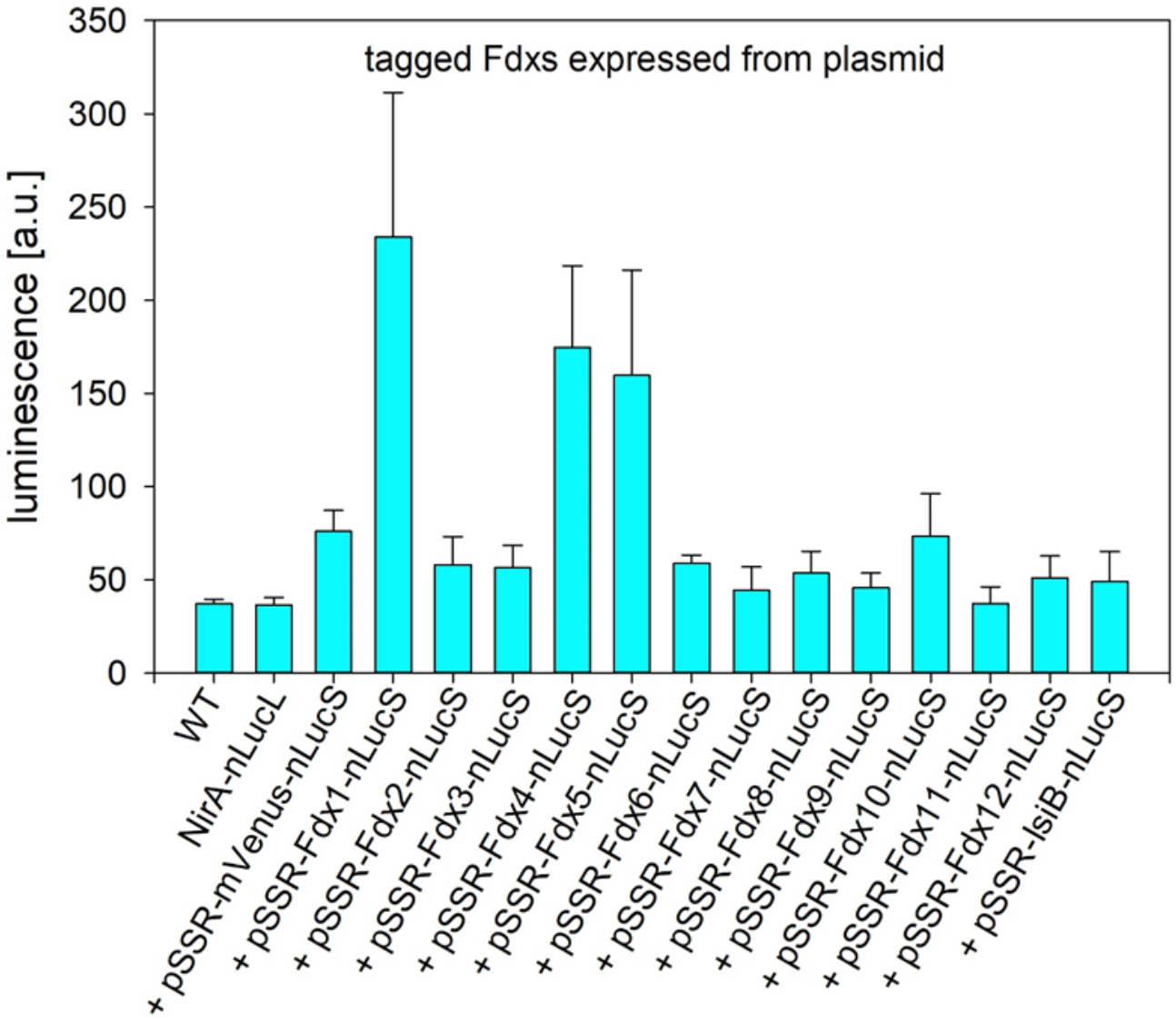
Detection of *in vivo* interaction between NirA and Fdxs 1-12 by luciferase complementation in cells grown under photoautotrophic conditions. Strains with a genome encoded NirA-nLucL fusion (nLucL, luciferase large subunit) and transformed with a plasmid (pSSR) encoded, rhamnose inducible, single Fdx-nLucS (Fdx, Fdx1-12; nLucS, luciferase small subunit). The bar is the luminescence level measured. Rhamnose inducible mVenus fluorescent protein was expressed from pSSR as a negative control. Plots are the means and standard deviations of three independent experiments.

In addition to the NirA binding assay, we measured the biochemical reactivity of specific Fdx proteins with two of the complexes that have significant roles in *Synechocystis* photosynthetic metabolism; PSI (F_B_^-^ → Fdx, -580 mV, (72) and PFOR (-540 mV, (73) (**Table 2**). Whereas PSI functions under photosynthetic growth conditions, PFOR functions under both highly reducing photomixotrophic and dark anaerobic metabolisms (18). Both have relatively low redox potentials, and it is feasible, based solely on the free-energy requirements, that these complexes can participate in electron transfer and reduction of each Fdxs 1-11 (**Table 2, Fig. S12**).

**Table 2.** Summary of electron transfer reactivity of Fdxs and IsiB with PSI, PFOR and HOX.

| Protein | Reduction<sup>1</sup> and oxidation<sup>2</sup> reactions |  |  |  |
| --- | --- | --- | --- | --- |
| | PSI<sup>3</sup><br>$E_m = -580$ mV | PFOR<br>$E_m = -540$ mV | HoxEFUYH<br>$E_m = -413$ mV<br>pH 7, 1 atm H<sub>2</sub> | HoxEFU<br>$E_m = -320$ mV, NAD(P)<sup>+</sup>/NAD(P)(H) |
| Fdx1 | + | + | + | + |
| Fdx2 | +/- | + | + | + |
| Fdx3 | + | + | + | n.d. |
| Fdx4 | + | - | + | + |
| Fdx5 | - | n.d. | n.d. | - |
| Fdx6 | +/- | n.d. | n.d. | n.d. |
| Fdx7 | +/- | - | n.d. | n.d. |
| Fdx8 | + | n.d. | n.d. | n.d. |
| Fdx9 | n.d. | n.d. | n.d. | n.d. |
| Fdx10 | - | - | n.d. | n.d. |
| Fdx11 | - | + | n.d. | + |
| Fdx12 | n.d. | n.d. | n.d. | n.d. |
| IsiB | + | + | + | + |

<sup>1</sup> PSI reduction reactions kinetics (**Fig. S14**); Fdx1, Fdx4 and IsiB also from (74) and PFOR-Fdx reduction assays (**Fig. S15**); Fdx1-3, 11 and IsiB also from (18).
<sup>2</sup> Previous studies of HoxEFU reactions were measured as either NAD(P)<sup>+</sup> reduction coupled to Fdx oxidation, or NADH oxidation coupled to Fdx reduction (12, 44). HoxEFUYH production of H<sub>2</sub> from reduced Fdx1-4 and IsiB were measured in cell-free lysates (11).
<sup>3</sup> +, electron transfer was observed; -, electron transfer was not observed. +/-, reactivity was above a threshold of 30% of electrons transferred for the 5:1 Fdx:PSI ratio only (**Fig. S14**). n.d., not determined. Reactivities of Fdx9 and Fdx12 were not tested.

In addition to the previously reported interaction of PSI with Fdx1 and Fdx4 (74) we demonstrate that Fdx1, Fdx3, Fdx4, Fdx8 and IsiB can function in electron transfer with PSI (**Table 2, Fig. S14**). Electron transfer from PSI was observed above the 30% electrons transferred threshold, but only for the 5:1 Fdx/PSI ratio sample in reactions with Fdx2, Fdx6 and Fdx7, suggesting that these Fdxs may not directly function in photosynthetic electron transport (**Table 2, Fig. S14**). Biochemical reactions with PFOR demonstrated that PFOR was able to reduce Fdx1, Fdx2, Fdx3, Fd11 and IsiB, but not Fdx4, Fdx7 and Fdx10 (**Table 2, Fig. S15**). Fdx5, Fdx6 and Fdx8 were not tested in these reactions.

Though not directly tested here, a summary of the Fdx reactivity profiles with HoxEFU and HoxEFUYH (12, 41) is shown based on being required for growth on BG11_0_+Arg+Glc media. In agreement with the upregulation of Fdx 4 and Fdx11 under these conditions, they both support NAD^+^ reduction by HoxEFU, and Fdx4 (together with Fdx1, 2, 3, and IsiB) supports H_2_ production by HoxEFUYH (**Table 2**). Overall, the activity results, together with the expression patterns (**Figure 4**), identify that different Fdxs are important under different growth regimes. Fdxs 1-4 and 11 all showed electron transfer activity that varied for the enzymes reported in Table 2.

## Discussion

The diversity of the ferredoxin isoforms in *Synechocystis* serves as a specialized “electron circuitry” designed to maintain cellular redox homeostasis. While plant-type [2Fe-2S] ferredoxins are well-documented in cyanobacteria as the primary drivers of photosynthetic electron transport, this study illuminates the roles of the less understood adrenodoxin-type, thioredoxin-like, and bacterial-type isoforms. These proteins expand the function from distribution of photoelectrons and catalysis to sensing and metabolic regulation.

Fdx1 is an essential protein, strongly expressed across a range of growth conditions and supports electron transfer with a broad range of redox enzymes. Fdx3 is the second most abundant ferredoxin under the conditions tested and expressed at similar levels across growth conditions, except for during Fe-limitation (**Figure 4, Figure S10**). In contrast, Fdx4 and Fdx8 are specifically induced under nitrate depletion, while Fdx10 is upregulated under iron limitation. Even though Fdx4 and Fdx5 are encoded within the same operon and Fdx4 has been shown to act as an electron transfer protein, Fdx5 did not have Fdx activity in reactions tested and shows divergent expression levels in this work. This suggests that *Synechocystis* utilizes post-transcriptional control to fine-tune the availability of specific isoforms in response to environmental signals.

Redox potentials range from -243 mV to -516 mV, suggesting that, thermodynamically, all isoforms, except for Fdx2, should support electron transfer to a broad range of dependent enzymes, including nitrite reductase (NirA). (**Fig. S12**), while PSI and PFOR could potentially function as electron donors. Furthermore, interactions with protein partners are determined by surface charge and kinetic gating, in addition to thermodynamic driving force. Plant-type Fdxs generally possess acidic (negative) surfaces that “steer” them toward positively charged docking sites on partners like PSI or FNR. Our in silico analysis revealed that Fdx10 and Fdx12 stand out with basic (positive) surface charges, an anomaly that suggests they evolved to interact with a unique set of acidic protein partners involved in specialized signaling rather than mainstream photosynthesis (69). Unsuccessful reduction of Fdx5 is inconsistent with the relatively high Fdx5 redox potential value measured by square wave voltammetry (-280 mV). Our data suggest that oxidation state of Fdx5 is likely dependent conformational rearrangements that may regulate Fe–S cluster accessibility and impede its reduction (18).

Under photomixotrophic conditions, the simultaneous input of electrons from water splitting and glucose oxidation challenges cellular redox homeostasis. High NADH levels typically inhibit the pyruvate dehydrogenase (PDH) complex, leading PFOR to functionally replace PDH in carbohydrate catabolism. We experimentally verified that PFOR directly reduces Fdx3 *in vitro* and determined that Fdx3 (*E*_m_ = -464 mV) and Fdx9 (*E*_m_ = -520 mV) possess the most negative redox potentials in the network. This low-potential capacity provides the thermodynamic prerequisite for their role as an electron sink, illustrating how *Synechocystis* leverages Fdx diversity to fine-tune redox homeostasis under extreme reducing pressure.

Fdx7 was previously reported to be involved in selenate tolerance (22). In this context it was shown to interact with FTR and glutaredoxin 2 (Grx2), the latter of which reduces selenate to less toxic forms (22). The exact electron pathway of this Fdx-glutaredoxin-thioredoxin crosstalk to selenate involving Fdx7 was not clarified in detail. Our findings close this gap, by suggesting that Fdx7 directly accepts electrons from PSI, to be utilized for mitigation of oxidative and metal stress through the Fdx-glutaredoxin-thioredoxin pathway.

Until now, only scarce information has been available regarding the function of Fdx8 other than it is essential for survival (5, 18) and that its expression is elevated under Fe-limitation and high-light (69). Our results demonstrate that Fdx8 transcription is also up-regulated under nitrogen depletion conditions. Here we demonstrated Fdx8 involvement in cell adaptation under nitrogen starvation conditions, which affects the cell capacity to catalyze nitrate reduction and carbon fixation. Interaction with PSI (**Table 2**), suggests that Fdx8 might be involved in coupling photosynthetic electron transport to other terminal acceptors, to maintain energy balance under nitrogen limitation.

While Fdx8 is essential for survival, even though its expression is relatively low under most conditions, it is strongly upregulated under nitrogen starvation and also in BG11_0_+Arg+Glc, suggesting that it serves a vital role during nutrient-induced stress. Although the Fe-S cluster binding motifs of Fdx8 predict the coordination of a [3Fe-4S] and a [4Fe-4S] cluster (5), we confirmed experimentally the presence of 2x[4Fe-4S] clusters, the level of which was dependent on oxygen conditions. Fdx8 may fine-tune its electron-transfer properties as oxygen levels change. For example, one of the two forms could either be inactive or confer different reaction kinetics of Fdx8 with redox partners (e.g., PSI). Along this line, aconitase, an enzyme of the TCA cycle that catalyzes the conversion of citrate to isocitrate, contains a [4Fe-4S] cluster which can reversibly form a [3Fe-4S] cluster leading to a change in protein function as a mRNA binding iron regulatory protein (60). The newly identified Fdx10 is a thioredoxin-type ferredoxin. Its classification and inability to accept electrons from PSI, combined with its unique basic surface charge, strongly indicates a role in redox-dependent protein regulation and stress-sensing rather than linear electron flux. Finally, Fdx11 is induced under nitrogen starvation and also in BG11_0_+Arg+Glc. Fdx11 supports electron transfer from PFOR and NADH reduction catalyzed by the [NiFe]-hydrogenase diaphorase HoxEFU complex, thereby facilitating metabolic adaptation under highly reducing conditions, such as photomixotrophic growth.

Ultimately, the *Synechocystis* Fdx suite represents a division of labor between high-volume photosynthetic electron flux and specialized metabolic regulation. By moving beyond the well-known plant-type proteins, we demonstrate how this suite of specialized ferredoxins allows the cell to balance the competing demands of efficient energy harvest and robust stress protection.

## Acknowledgments

We acknowledge funding by the Bundesministerium für Bildung und Forschung (BMBF) in the framework of the project CyFun (03SF0652A) and the Dietmar Hopp Stiftung. This work was authored in part by the National Laboratory of the Rockies for the U.S. Department of Energy (DOE), operated under Contract No. DE-AC36-08GO28308. Funding provided to D.S., C.E.L., D.W.M., E.C.K., Z.G., A.L., P.W.K. by the U.S. Department of Energy Office of Basic Energy Sciences, Division of Chemical Sciences, Geosciences, and Biosciences, Photosynthetic Systems Program. The views expressed in the article do not necessarily represent the views of the DOE or the U.S. Government. The U.S. Government retains and the publisher, by accepting the article for publication, acknowledges that the U.S. Government retains a nonexclusive, paid-up, irrevocable, worldwide license to publish or reproduce the published form of this work, or allow others to do so, for U.S. Government purposes. We thank Jonas Quast for the growth and harvest of cell material that was used in this study.

## Declaration of generative AI and AI-assisted technologies in the manuscript preparation process

During the preparation of this work the author(s) used ChatGPT (Open AI 2025), ChatGPT Team (5.1) and DeepL in order to optimize language. After using this tool/service, the authors reviewed and edited the content as needed and take full responsibility for the content of the published article.

## Supplementary Information

The supplementary information file contains Supplemental Text S1-3, Figures S1-S17 and Tables S1-S4.

