## Supplementary Information for "Comprehensive study on ferredoxin isoforms in the cyanobacterium Synechocystis sp. PCC 6803"

### Supporting Text 1

**Construction of mutagenesis plasmids.** To probe ferredoxin expression levels under different growth conditions, 6xHis-tagged mutant strains for ferredoxins 1 to 11 were generated. Each mutagenesis plasmid was generated from a C-entry vector (opened with *Bam*HI and *Eco*RI) into which a cassette excised from a toolkit vector (with *Bam*HI and *Eco*RI) was ligated. The C-entry plasmids used the pBluescript SK+ plasmid (digested with *Kpn*I and *Sac*I) as a backbone and contained two overlapping ~250 bp DNA fragments (up- and downstream of the respective stop codon of the gene of interest) that were introduced via Gibson cloning. The primers 29 to 72 (**Table S4**) used to generate the two fragments further introduced the following bases 5'-CCCATCGGA-3' followed by *Bam*HI, *Xho*I and *Eco*RI restriction sites directly before the stop codon. The toolkit vector for the 6xHis-GENT cassette was constructed by inserting a PCR amplified gentamycin resistance cassette (primers 73 and 74) by blunt-ended ligation into the *Eco*RV site of pMB0122. The final mutagenesis plasmids were sequenced (Azenta) and transformed into the *Synechocystis* wild-type strain to create the 6xHis-tagged ferredoxin mutant strains. Mutant segregation was confirmed by PCR (e.g. using primer 29 and 32 for Fdx1).

For the nanoLuc *in vivo* interaction studies between nitrite reductase and Fdxs, the LgBit luciferase (nLucL) coding fragment together with a gentamycin resistance cassette was excised from a toolkit vector (pMB00071) using *Bam*HI and *Eco*RI, ligated into the *Bam*HI and *Eco*RI opened NirA C-entry plasmid (ordered from Azenta; NirA contains an internal *Bam*HI site that was silently mutated in this C-entry vector) and transformed into *Synechocystis*. The resulting mutant was confirmed by PCR using primers 75 and 76 (**Table S4**) and then transformed either with plasmids that (A) added the SmBit luciferase (nLucS) together with a spectinomycin cassette (*Bam*HI and *Eco*RI excised from the toolkit vector pMB0070) before the STOP codon of the investigated Fdxs (using the same C-entry vectors mentioned above) or (B) with a rhamnose inducible plasmid for the expression of a respective ferredoxin tagged with the SmBit luciferase (nLucS) tag. These plasmids derived from a plasmid that has been described in the literature (1) and into which the SmBit coding region was introduced, so that a gene of interest could be incorporated into the vector using a NdeI restriction site. The primers used to generate the Fdx fragments that were incorporated into this vector via Gibson cloning were primers 77 to 100 (**Table S4**). Mutants were confirmed by PCR using the same primers as for the His-tagged strains for (A) and primers 103 and 104 for (B) (**Table S4**).

### Supporting Text 2

**LC/MS analysis.** For protein identification following purification, 0.5 µL of each sample, containing 2–14 µg of protein, was mixed with SDS loading buffer and heated at 95°C for 5 min. As a preliminary cleaning step, the boiled samples were run into a 4 % SDS PAGE stacking gel at 120 V. After the entire sample exited the well and was captured within the gel, the proteins were excised using a ethanolcleaned razor blade. The excised gel pieces were prepared for protein mass spectrometry *via* a nano-liquid chromatography tandem mass spectrometer (nLC-MS/MS) through a series of steps: destaining, reduction of disulfide bridges, cysteine carbamidomethylation, and tryptic in-gel digestion, following an established protocol (2). The resulting peptides were extracted in multiple steps using 50 % (v/v) acetonitrile, 0.1 % (v/v) formic acid, or 5 mM ammonium bicarbonate, and subsequently desalted using ZipTip-μ$C_{18}$ material (Merck Millipore). After desalting, the peptides were dried in a vacuum centrifuge and resuspended in 20 μL of 0.1 % (v/v) formic acid before mass spectrometry analysis on an Orbitrap Fusion Tribrid mass spectrometer (Thermo Scientific) coupled to a nLC system Dionex Ultimate 3000RSLC (Thermo Scientific) *via* an electrospray ion source TriVersa NanoMate (Advion). A 5-μL volume containing the peptides was separated on an Acclaim PepMap100 $C_{18}$ column (Thermo Scientific) at a flow rate of 300 nL $min^{-1}$, following the parameters described elsewhere (3).

Proteome Discoverer v2.2 (Thermo Scientific) was employed for protein identification using the SequestHT search engine against the proteome databases of the production strains *E. coli* K12 (taxonomy ID: 83333), used for the heterologous production of Fdx1 to Fdx11, or *E. coli* BL21 (DE3) (taxonomy ID: 511693), used for the production of IsiB, from the UniProt database, supplemented with the amino acid sequences of the target proteins (Fdx1–Fdx11 and IsiB). The following parameters were set: carbamidomethylation of cysteine residues as a fixed modification, oxidation of methionine as a dynamic modification; trypsin cleavage specificity with a maximum of two missed cleavage sites; precursor and fragment mass tolerances set to 3 ppm and 0.6 Da, respectively. Stringent identification criteria were applied with a 1% false discovery rate threshold for peptide identification using the Target Decoy PSM Validator node. Protein quantification was facilitated by the Minora node in Proteome Discoverer through label-free quantification based on MS1 precursor intensity values. Relative protein abundance was calculated by normalizing the MS1 intensities of the target proteins in the gel slices to the overall MS1 intensities of all detected proteins within the same slices.

### Supporting Text 3

**EPR analysis on the effect of oxygen on Fdx8.** To investigate the nature of the Fdx8 Fe-S clusters and the impacts of oxygen, we conducted EPR experiments on Fdx8 expressed under aerobic and strictly anaerobic conditions in the “as purified” state or reduced with 5 mM sodium dithionite (**Fig. S9**). These samples were not reconstituted, in contrast to the EPR spectrum of reduced Fdx8 displayed in the main text (**Fig. 3**), as well as previous studies on other di-cluster ferredoxin or ferredoxin-like proteins (4, 5). Generally, EPR signals of oxidized [3Fe-4S] clusters can be detected while reduced [3Fe-4S] clusters are EPR silent. The opposite is true for [4Fe-4S] clusters, where EPR signals of reduced [4Fe-4S] clusters can be detected and oxidized [4Fe-4S] clusters are EPR silent.

**Supporting Text 4**

**Purification of PSI protein complexes.** PSI protein complexes employed for the P700+ reduction kinetics measurements were purified from a PsaF-6xHis mutant strain constructed as described elsewhere (6) using a previously described protocol (7, 8) with modifications. Briefly, a 30-L *Synechocystis* PsaF-6xHis culture was harvested (15 min, 4°C, 5.000 x g) and resuspended in buffer A (20 mM MES pH 6.5, 10 mM $MgCl_2$, 10 mM $CaCl_2$). Cells were broken using a multi-cycle cell disruptor at 35 kpsi (Constant Cell disruption Systems). Membranes were pelleted (10 min, 4°C, 20.000 x g), washed in buffer A, pelleted (10 min, 4°C, 20.000 x g), washed in buffer B (20 mM MES pH 6.5, 10 mM $MgCl_2$, 10 mM $CaCl_2$, 500 mM mannitol) to which fresh 0.05 % (w/v) beta-dodecyl-D-maltoside was added and pelleted again (10 min, 4°C, 20.000 x g). Subsequently, the membranes were resuspended in extraction buffer (20 mM HEPES pH 7.5, 10 mM $MgCl_2$, 10 mM $CaCl_2$, 200 mM ammonium sulfate), the Chl a concentration adjusted to 1 mg/ml and the membranes solubilized with 0.5 % (w/v) N,N-Dimethyl-n-dodecylamine N-oxide (LDAO) for 30 min at RT with gentle agitation. After pelleting insoluble material (60 min, 4°C, 100.000 x g), the supernatant was applied to a prepacked fast flow Co-TALON resin cartridge (Takara) connected to an Akta pure (Cytiva) with (20 mM HEPES pH 7.5, 10 mM $MgCl_2$, 10 mM $CaCl_2$, 5 mM imidazole, 0.03 % (w/v) beta-dodecyl-D-maltoside) as the running buffer. The sample was washed with 10 CV running buffer and after elution with (20 mM HEPES pH 7.5, 10 mM $MgCl_2$, 10 mM $CaCl_2$, 100 mM imidazole, 0.03 % (w/v) beta-dodecyl-D-maltoside) green fractions were pooled, concentrated and the Chl a concentration determined. Prior to flash freezing and storage at -80°C the PSI protein complexes were mixed 1:1 with a PSI storage buffer (20 mM HEPES pH 7.5, 10 mM $MgCl_2$, 10 mM $CaCl_2$, 1 M mannitol, 0.03 % (w/v) beta-dodecyl-D-maltoside).

### Supporting Figures

**Fdx1 (Ssl0020)**
>sp|P27320|FER_SYNY3 Ferredoxin-1 OS=Synechocystis sp. (strain PCC 6803 / Kazusa) OX=1111708 GN=petF PE=1 SV=2
MASYTVKLITPDGESSIECSDDTYILDAAEEAGLDLPYSCRAGACSTCAGKITAGSVDQSDQSFL
DDDQIEAGYVLTCVAYPTSDCTIETHKEEDLY

**Fdx2 (Sll1382)**
>tr|P74159|P74159_SYNY3 Ferredoxin OS=Synechocystis sp. (strain PCC 6803 / Kazusa) OX=1111708 GN=petF PE=3 SV=1
MSRSHRVLIHDRQNEKDYSVIVSDDRYILHQAEDQGFELPFSCRNGACTACAVRVISGQIHQPEA
MGLSPDLQRQGYALLCVSYAQSDLEVETQDEDEVYELQFGRYFGAGRVRLGLPLDED

**Fdx3 (Slr1828)**
>tr|P73388|P73388_SYNY3 Ferredoxin OS=Synechocystis sp. (strain PCC 6803 / Kazusa) OX=1111708 GN=petF PE=3 SV=1
MVNTYTAEIQHQGQTYTISVPEDKTVLQAADDEGIQLPTSCGAGVCTTCAALITEGTAEQADGMG
VSAELQAEGYALLCVAYPRSDLKIITEKEDEVYQRQFGGQG

**Fdx4 (Slr0150)**
>tr|P74449|P74449_SYNY3 Ferredoxin OS=Synechocystis sp. (strain PCC 6803 / Kazusa) OX=1111708 GN=petF PE=3 SV=1
MGAIYSVNLVNPATGSDVTIEVAEDELILEAAENQGLDLPYSCRAASCVACAGRLLEGTVEHTDK
GSDFLKPEELAAGCVLLCAAYATSDCKILTHQEEALFG

**Fdx5 (Slr0148; sequence is actually shorter than listed in the UniProt database (without the highlighted sequence in grey, the valine at the start is a methionine))**
http://alcodb.jp/cyano/PCC6803/slr0148/network
>tr|P74447|P74447_SYNY3 Ferredoxin OS=Synechocystis sp. (strain PCC 6803 / Kazusa) OX=1111708 GN=slr0148 PE=3 SV=1
MTMPPLWNCSVANRVNAIVASTKEDCMAKTIKLDPIDLKVAIETNDNLLSGLLGQDLRIMKECGG
RGMCATCHVYITAGMESLSPLNRREQRTLEVITTHNRYSRLACQARVLDEGVVVELPAGMYVSE
IEDIEELIGRRAEENILNPRDGSILVEKGKLITRSMISQLDDQLQAAKIQIVNDTDE

**Fdx6 (Ssl2559)**
>tr|P73171|P73171_SYNY3 Ferredoxin OS=Synechocystis sp. (strain PCC 6803 / Kazusa) OX=1111708 GN=ssl2559 PE=4 SV=1
MNNCVISFPQTKFLPLSLEFNACLAEYLTPDNSPILFGCRTGLCGTCLVKVVGEILSPEAEEREILA
ILAPDDVQARLACQIKLTGDIAIRAYQSDEI

**Fdx7 (Sll0662)**
>tr|Q55980|Q55980_SYNY3 Ferredoxin OS=Synechocystis sp. (strain PCC 6803 / Kazusa) OX=1111708 GN=sll0662 PE=1 SV=1
MVIADLNFPPDHRSGLEPELGGDWRNFDDRSGLEPELGGELRERGVYVDEVTCIGCKNCAHVA
PNTFTIEQEHGRSRAFSQNGDDEAVIQEAIDTCPVDCIHWVPYDELKHLEEKRKHQQIRPLGYPQ
INPHL

**Fdx8 (Sssr3184)**
>tr|P73649|P73649_SYNY3 Ferredoxin OS=Synechocystis sp. (strain PCC 6803 / Kazusa) OX=1111708 GN=ssr3184 PE=4 SV=1
MPHTIVTETCEGVADCVEACPVACIHPGDGKNTIGTDWYWIDFATCIDCGICLQVCPVEGAILPEE
RPDLQKSPA

**Fdx9 (Slr2059)**
>tr|P73811|P73811_SYNY3 Ferredoxin OS=Synechocystis sp. (strain PCC 6803 / Kazusa) OX=1111708 GN=slr2059 PE=4 SV=1
MKKRVTLTFPRSAVQMPVTYRLAKDFNIAANIIRAQVAPNQVGKVVLELSGDIDQLEASLEWMRS
QSIEVSLASREIVIDDQSCVDCGLCTGVCPTEALSLDPDSFRLMFRRSRCVVCEQCIPSCPVQAIA
TNF

**Fdx10 (Sll1584)**
>tr|P73195|P73195_SYNY3 Sll1584 protein OS=Synechocystis sp. (strain PCC 6803 / Kazusa) OX=1111708 GN=sll1584 PE=4 SV=1
MENPVPITPSETIVDSCQRLGLGRIQRHLFLCCDQTKPKCCSKEDSLATWDYLKKRLPELGLDCT
QSSRDGNIFRTKANCLRVCQQGPILLVYPEGIWYRNVTPTVMEKILQEHILQNRPVEEYRFFTHPL
SHL

Note: Without structural analysis it is not possible to predict which one of two C residues bind the Fe-S cluster.

**Fdx11 (Ssl3044)**
>tr|P74283|P74283_SYNY3 Ferredoxin OS=Synechocystis sp. (strain PCC 6803 / Kazusa) OX=1111708 GN=ssl3044 PE=3 SV=1
MTITFVKEQKDIVVAQGANLREKALQNGVDIYTLKGKLMNCGGYGQCGTCIVEITAGMENLSPKT
DFENRVLRKKPDNFRLACQTLVNGPVSVNTKPKG

**Fdx12 (Ssr1041)**
>tr|P74801|P74801_SYNY3 Ssr1041 protein OS=Synechocystis sp. (strain PCC 6803 / Kazusa) OX=1111708 GN=ssr1041 PE=4 SV=1
MAVTIHFLPDDVTVAARVGEPILDVAERAGVFIPTGCLMGSCHACEVELGDGTPICACISAVPVGV
QELEINLYDDLTW

**Fig. S1.** Amino acid sequences of Fdx1-Fdx12 from *Synechocystis* p. PCC 6803. Cysteines coordinating Fe-S clusters are highlighted in yellow, other cysteines are highlighted in cyan.

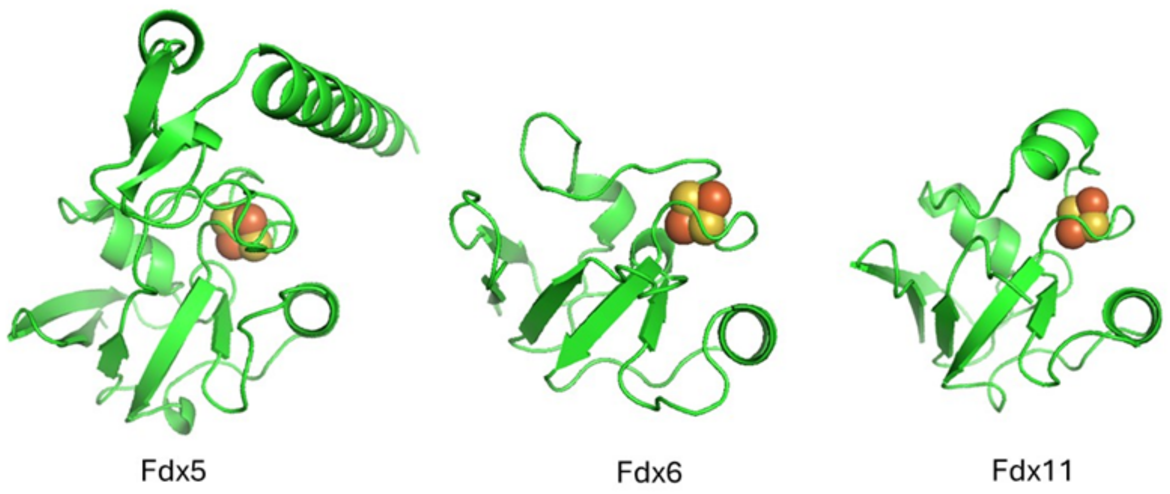


**Fig. S2.** Structures of Fdx5, Fdx6 and Fdx11. The same models presented as in Figure 1, in a different orientation.

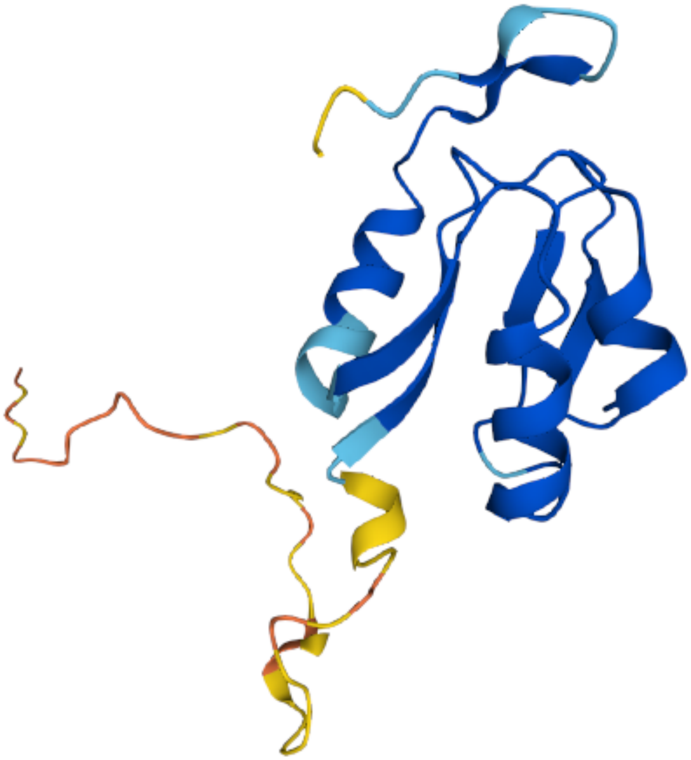

**Fig. S3.** AlphaFold-predicted structure of Fdx7, highlighting a poorly defined N-terminal region (yellow) with low confidence. The well-defined core, predicted with high confidence, is shown in blue.

```
Fdx8         ETCEGVA--DCVEACPVACIHPGDG--KNTIGTDWYWIDFATCIDCG------ICLQVCPVEGAILPEERPDLQKSPA 75
7Fe-8SST6I   DICEGIA--DCVNACPVACIHMGNG--INKKGTNFYWIDFNTCIDCG------ICLQVCPLENAILAEERSELQQIN- 74
2V2K         EPCVDVKDKACIEECPVDCIYEG---------ARMLYIHPDECVDCG------ACEPVCPVEAIYYEDDVPDQWSSYA 68
1H98         EPCIGVKDQSCVEVCPVECIYDG---------GDQFYIHPEECIDCG------ACVPACPVNAIYPEEDVPEQWKSYI 68
1BD6         EPCIGTKDASCVEVCPVDCIHEG---------EDQYYIDPDVCIDCG------ACEAVCPVSAIYHEDFVPEEWKSYI 68
1XER         DLCIADG--SCINACPVNVFQWYDTPGHPASEKKADPVNEQACIFCM------ACVNVCPVAAIDVKPP-------- 103
sp|P08813|   DKCTGDG--ECVDVCPVEVYELQD--------GKAVPVNEEECLGCE------SCVEVCEAGAITVEEN--------- 62
6FD1         DNCIKCKYTDCVEVCPVDCFYEG---------PNFLVIHPDECIDCA------LCEPECPAQAIFSEDEVPEDMQEFI 68
1DUR         DSCIACG--ACKPECPVNCIQEG----------SIYAIDADSCIDCG------SCASVCPVGAPNPED---------- 55
1CLF         DSCVSCG--ACASECPVNAISQG---------DSIFVIDADTCIDCG------NCANVCPVGAPVQE----------- 55
2FDN         EACISCG--ACEPECPVNAISSG---------DDRYVIDADTCIDCG------ACAGVCPVDAPVQA----------- 55
1BLU         DECINCD--VCEPECPNGAISQG---------DETYVIEPSLCTECVGHYETSQCVEVCPVDCIIKDPSHEETEDELR 72
2FGO         DDCINCD--VCEPECPNGAISQG---------EEIYVIDPNLCTECVGHYDEPQCQQVCPVDCIPLDDANVESKDQLM 72
```

**Fig. S4.** Sequence alignment of Fdx8 with selected other [7Fe-8S] and [8Fe-8S] ferredoxins with available crystal structures. The figure displays a portion of the alignment, focusing on the key Fe-S binding sequence. Residues are color-coded based on their functional roles: cysteine residues coordinating the first Fe-S cluster are highlighted in black, those coordinating the second Fe-S cluster are in red, and conserved cysteine residues involved in hydrogen bonding one of the Fe-S clusters are highlighted in cyan. Residues **Y** (yellow) in position $CX_2$**Y**$X_{2\text{-}4}C$ determine the type of the first Fe-S cluster as [3Fe-4S] or [4Fe-4S], or exchangeable [3/4Fe-4S]. Conserved residues present in Fdx8 and across the sequences are shaded in gray. The ferredoxins sequences are derived from the following organisms*:* *Synechocystis sp*. PCC 6803 (Fdx8), *Paulinella chromatophore* (7Fe-8SST6), *Mycobacterium smegmatis* (2V2K), *Thermus aquaticus* (1H98), *Bacillus schlegelii* (1BD6), *Sulfolobus tokodaii* (1XER), *Desulfovibrio vulgaris* (sp|P08813|), *Azotobacter vinelandii* (6FD1), *Peptoniphilus asaccharolyticus* (1DUR), *Clostridium pasteurianum* (1CLF), *Clostridium acidurici* (2FDN), *Allochromatium vinosum* (1BLU), and Pseudomonas aeruginosa (2FGO). These organisms represent diverse phyla including *Cyanobacteria, Actinobacteria, Firmicutes, Deinococcus-Thermus, Archaea, Proteobacteria, and Peptostreptococcaceae*. Sequence alignment was performed using Clustal Omega.

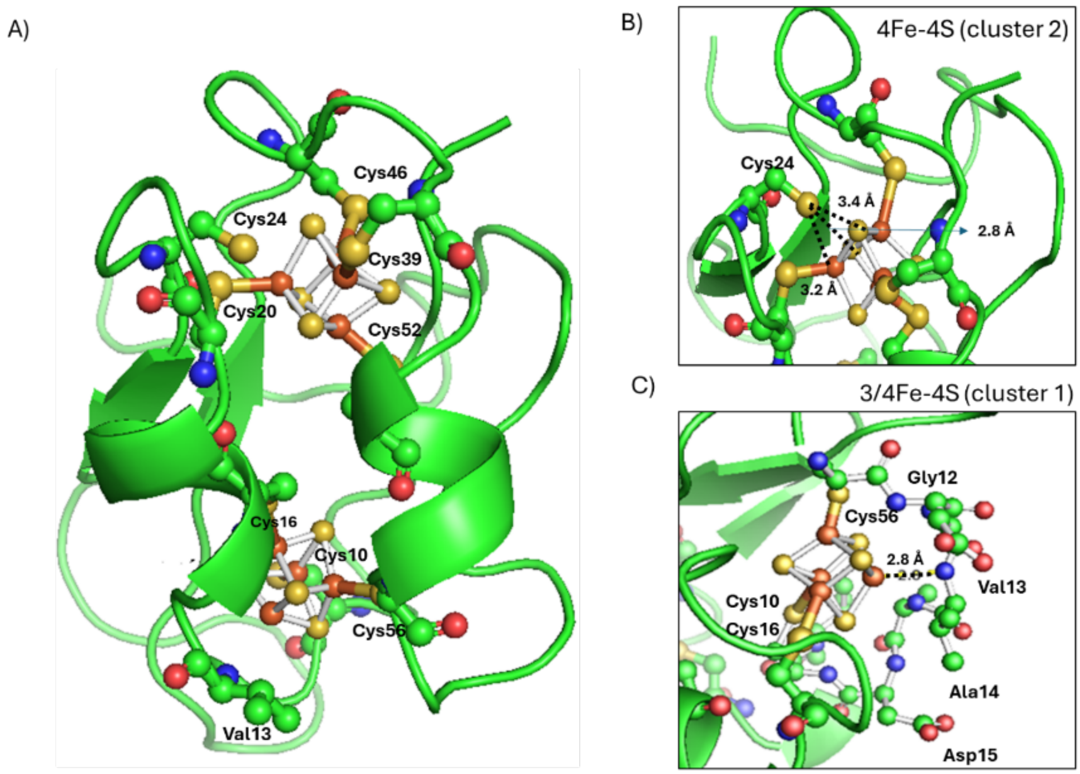


**Fig. S5.** Fdx8 structural model showing Cysteine residue distribution, as well as the position of Val13 in proximity to the [3/4Fe-4S] cluster (sequence coordinating the first cluster DC**V**$_{13}$EACPV; see Fdx8 sequence in Figure S4). The position of the Fe-S cluster was determined based on comparison with known structures PDB ID 1H98 from *Thermus aquaticus* using Pymol. Green = peptide, nonpolar residues; orange = iron; yellow = sulfur; blue = nitrogen; red = oxygen; dashed lines are distance between Fe and key residues (A) whole protein, (B) and (C) zoom in on [4Fe-4S] and [3/4Fe-4S] clusters, respectively. The roles of cysteine residues were assigned based on their proximity to the Fe-S clusters in the model structure. Residues Cys10, Cys16, and Cys56 coordinate the iron atoms in Cluster 1 ([3/4Fe-4S]), while residues Cys20, Cys46, Cys49, and Cys52 coordinate the iron atoms in Cluster 2 ([4Fe-4S]). Notably, the orientation and distance of the sulfur atom in Cys24 relative to Cluster 2 suggest that it does not directly coordinate an Fe atom. Instead, it likely forms a hydrogen bond with a sulfur atom in the cluster, as indicated by the 2.8 Å distance. Furthermore, the position and orientation of Val13 in the model suggest that its backbone nitrogen may coordinate an Fe atom in Cluster 1, also at a distance of 2.8 Å.

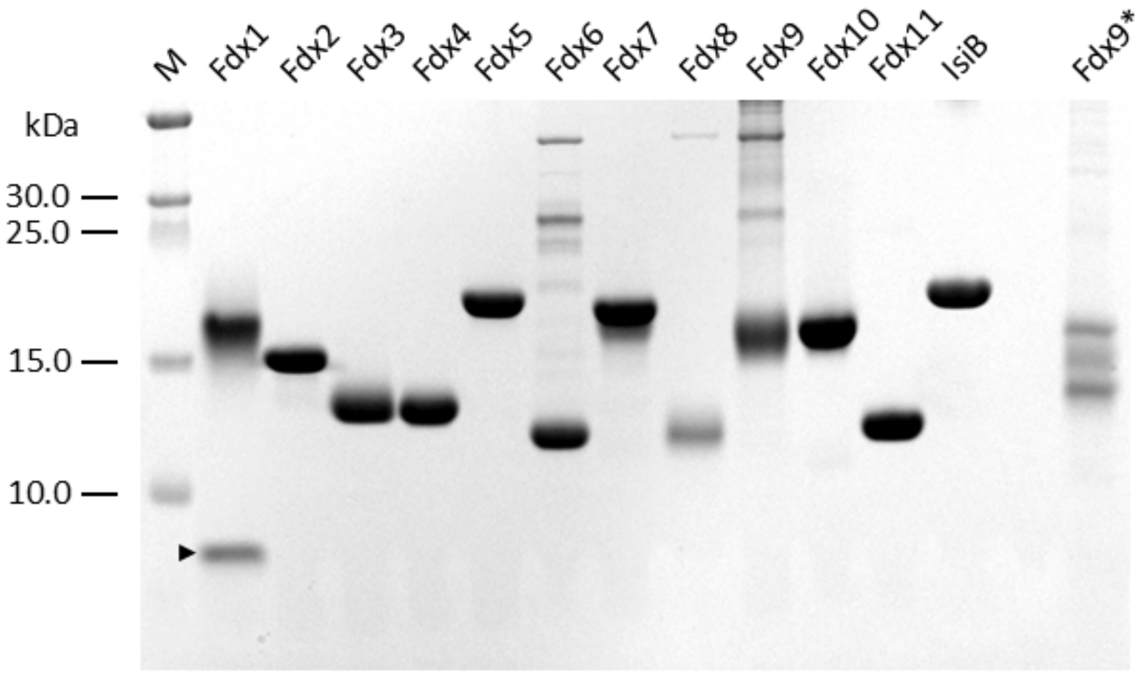


**Fig. S6.** SDS PAGE analysis of the purified *Synechocystis* Fdx1-Fdx11 and flavodoxin (IsiB) (see also Table S2). The arrowhead indicates the Fdx1 protein band. The other major band in that lane, just above 15 kDa, is maltose-binding protein (Accession # A0A384L126; detected by LC/MS, data not shown; for the method see **Supporting Text 2**), which was introduced into the sample at the TEV cleavage step (see Materials and Method) (see Table 2 for more details).

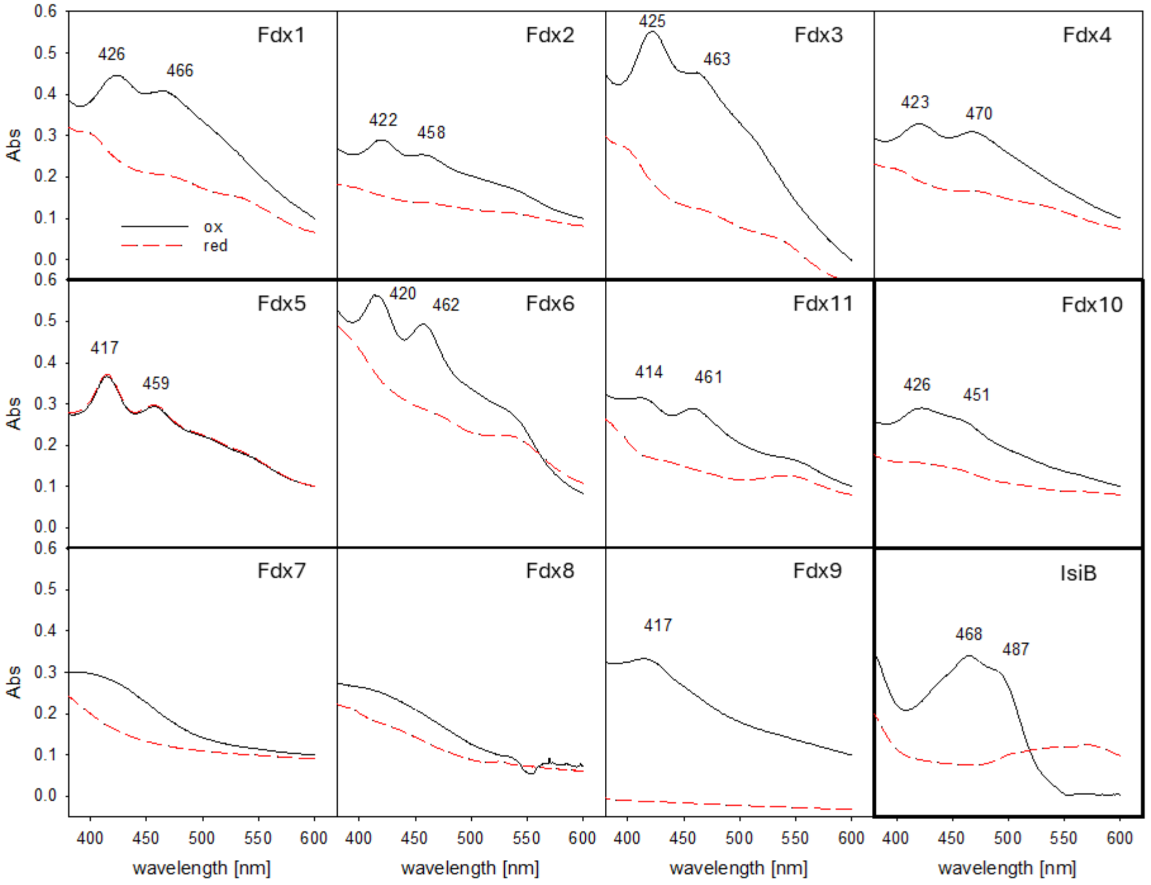


**Fig. S7.** UV-Vis absorption spectra of as purified Fdxs 1-11 and flavodoxin (black line), as well as spectra from reduced proteins (red dashed line) between 350 and 450 nm. Reduction conditions (a final concentration of 4 mM sodium dithionite) that for the other proteins lead to a change in absorption, did not change the spectrum for Fdx5. Buffer included 50 mM HEPES buffer, 0.1M NaCl and 5% glycerol at pH 8.

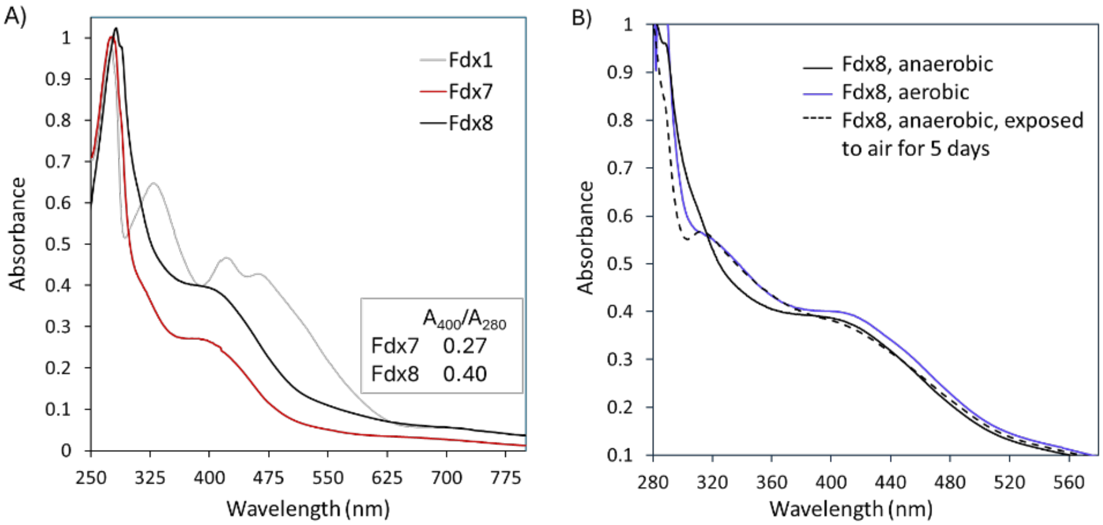


**Fig. S8.** (A) UV-Vis spectra of Fdx1, Fdx7, and Fdx8. Spectra were recorded using as-purified proteins. Fdx1 and Fdx7 were prepared under aerobic conditions, while Fdx8 was maintained strictly anaerobically. (B) UV-Vis spectra of Fdx8 under different preparation conditions: strictly anaerobic versus aerobic. The spectrum of anaerobically prepared Fdx8 suggests the presence of [4Fe-4S] clusters, whereas the aerobically prepared sample likely contains a mixture of [4Fe-4S] and [3Fe-4S] clusters. Upon exposure to air, spectral features characteristic of [3Fe-4S] clusters become more apparent. Fdxs were stored in 50 mM HEPES buffer, 0.1M NaCl and 5% glycerol at pH 8.

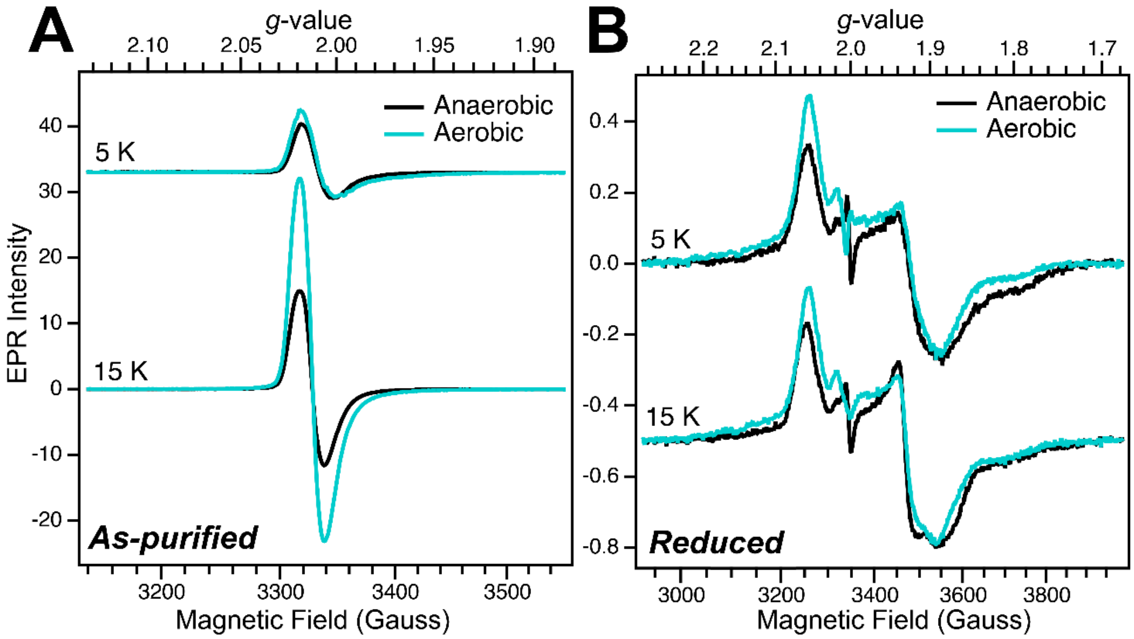


**Fig. S9.** EPR spectra of Fdx8 expressed and purified under aerobic (teal lines) or strictly anaerobic (black lines) conditions. EPR samples were prepared without any further treatment of the protein (i.e. no reconstitution was performed) and the as-purified spectra (A) collected. Samples were then thawed and treated with a high concentration stock of sodium dithionite (NaDT) to achieve a final concentration of 5 mM NaDT, re-frozen, and the reduced spectra (B) collected. Spectra collected using 1 mW of microwave power at either 5 K or 15 K sample temperature and are shown after cavity background subtraction and additional baseline correction using a polynomial spline.

**A**

| | Photoautotroph | Photomixotroph | Heterotroph | BG11 +Arg +Gluc | Nitrate depleted | Iron depleted | score mean |
|---|---|---|---|---|---|---|---|
| Fdx1 | 11 | 11 | 11 | 11 | 11 | 11 | 11,0 |
| Fdx2 | 5 | 8 | 5 | 3 | 4 | 7 | 5,3 |
| Fdx3 | 10 | 10 | 10 | 10 | 10 | 9 | 9,8 |
| Fdx4 | 4 | 5 | 4 | 5 | 7 | 5 | 5,0 |
| Fdx5 | 9 | 9 | 9 | 9 | 9 | 8 | 8,8 |
| Fdx6 | 2 | 4 | 0 | 4 | 3 | 0 | 2,2 |
| Fdx7 | 3 | 0 | 4 | 1 | 1 | 3 | 2,0 |
| Fdx8 | 0 | 0 | 0 | 2 | 7 | 0 | 1,5 |
| Fdx9 | 6 | 4 | 7 | 7 | 7 | 7 | 6,3 |
| Fdx10 | 7 | 7 | 8 | 7 | 2 | 10 | 6,8 |
| Fdx11 | 8 | 7 | 7 | 8 | 8 | 4 | 7,0 |

Score: 11 10 9 8 7 6 5 4 3 2 1 0 (highest to lowest)

**B**

| | Photoautotroph | Photomixotroph | Heterotroph | BG11 +Arg +Gluc | Nitrate depleted | Iron depleted |
|---|---|---|---|---|---|---|
| Fdx1 | 0,0 | 0,0 | 0,0 | 0,0 | 0,0 | 0,0 |
| Fdx2 | -0,3 | 2,7 | -0,3 | -2,3 | -1,3 | 1,7 |
| Fdx3 | 0,2 | 0,2 | 0,2 | 0,2 | 0,2 | -0,8 |
| Fdx4 | -1,0 | 0,0 | -1,0 | 0,0 | 2,0 | 0,0 |
| Fdx5 | 0,2 | 0,2 | 0,2 | 0,2 | 0,2 | -0,8 |
| Fdx6 | -0,2 | 1,8 | -2,2 | 1,8 | 0,8 | -2,2 |
| Fdx7 | 1,0 | -2,0 | 2,0 | -1,0 | -1,0 | 1,0 |
| Fdx8 | -1,5 | -1,5 | -1,5 | 0,5 | 5,5 | -1,5 |
| Fdx9 | -0,3 | -2,3 | 0,7 | 0,7 | 0,7 | 0,7 |
| Fdx10 | 0,2 | 0,2 | 1,2 | 0,2 | -4,8 | 3,2 |
| Fdx11 | 1,0 | 0,0 | 0,0 | 1,0 | 1,0 | -3,0 |

Deviation from mean: 5 4 3 2 1 0 -1 -2 -3 -4 -5 (increase to decrease)

**Fig. S10.** Qualitative analysis of Fdx expression levels for the immunoblot analysis shown in Figure 4. (A) Expression levels were assessed and compared by evaluating the signal strengths in the immunoblot analysis shown in Figure 4 of the main text by eye. The score ranges from 0 to 11 (11 = highest expression, 0 = no detectable expression). When two signals appeared equally strong, they received the same score, and the next intense band was scored accordingly lower. In some cases, the assessment and scoring were hampered by nearby unspecific signals from the $\alpha$His antibody. While a comparison within one panel is allowed, a comparison between two different panels may be flawed due to potentially different exposure times. Mean scores were calculated over all the tested conditions. (B) The calculated means were subtracted from the scores given in panel A. A positive value therefore indicates a relative increase in expression compared to the other conditions, while a negative value indicates a relative decrease.

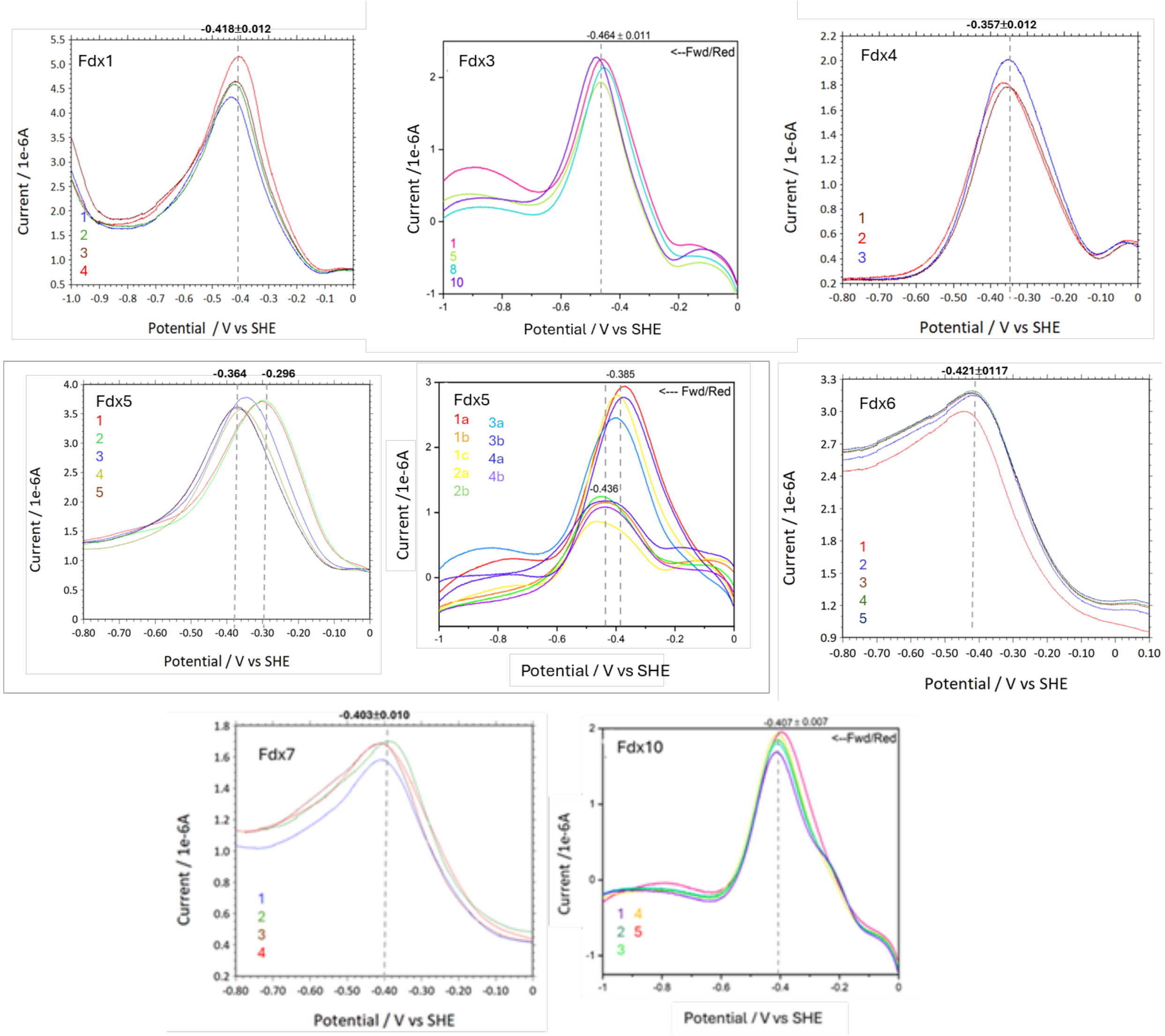


**Fig. S11.** Redox potentials were measured using square wave voltammetry with proteins immobilized on a polished pyrolytic graphite edge (PGE) electrode in HEPES buffer (pH 8.0). Potentials were converted to values vs. SHE by adding 199 mV to the Ag/AgCl reference. Numbers indicate subsequent scans measured for one or more Fdx electrodes.

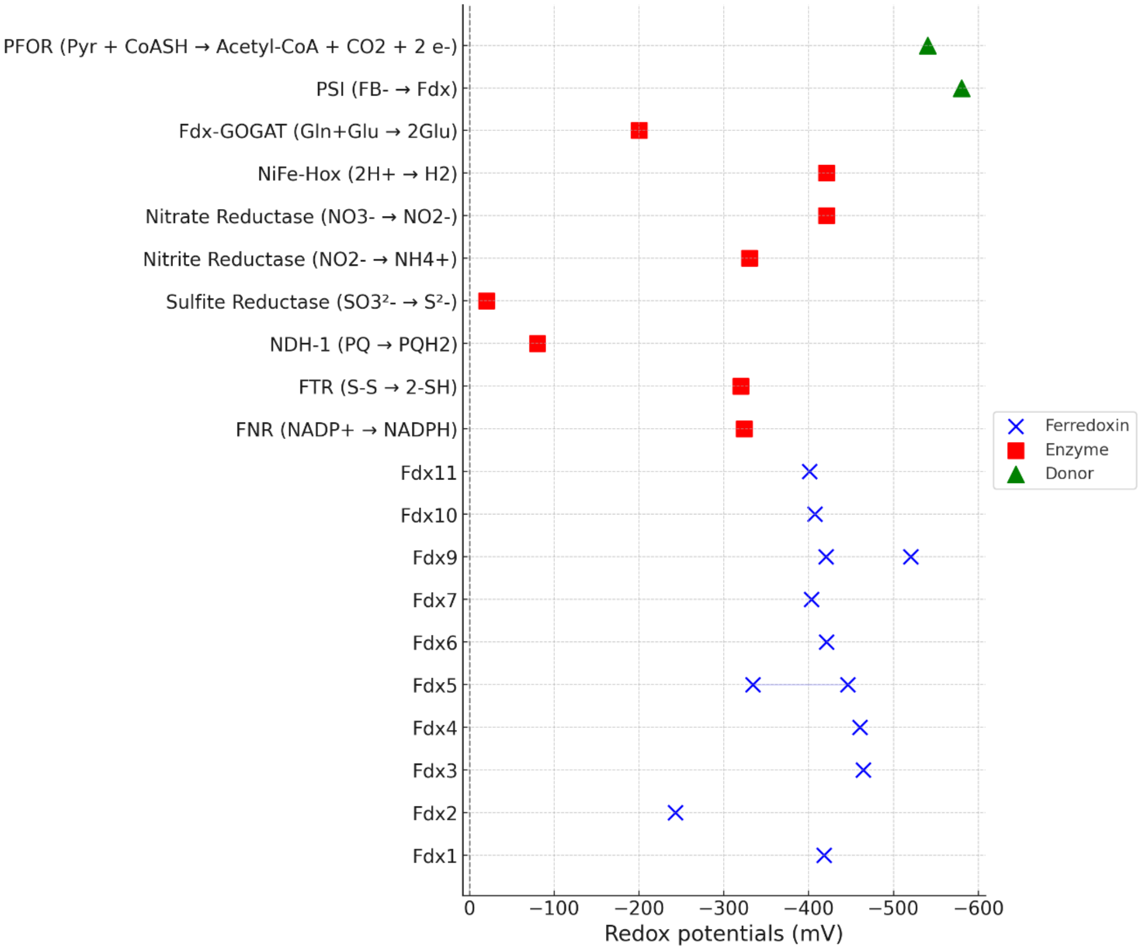


**Fig. S12.** Comparison of redox midpoint potentials ($E_m$) for *Synechocystis* Fdxs and redox potentials (Em) of reactions catalyzed by relevant protein partners. For Fdx4 the Em value from the literature (9) is included. There are two measured $E_m$'s for Fdx5, -280 and -430 mV (Table 1). The Em values for the protein partners were obtained from the following references: FNR (10), FTR (11), Hox (10), Fdx-GOGAT (12), NDH1 (13), SR (sulfite reductase) (14), NR (nitrate reductase) (10), NirA (nitrite reductase) (10), PSI (15), and PFOR (16). The value for Flv1/3 is +816 mV (10).

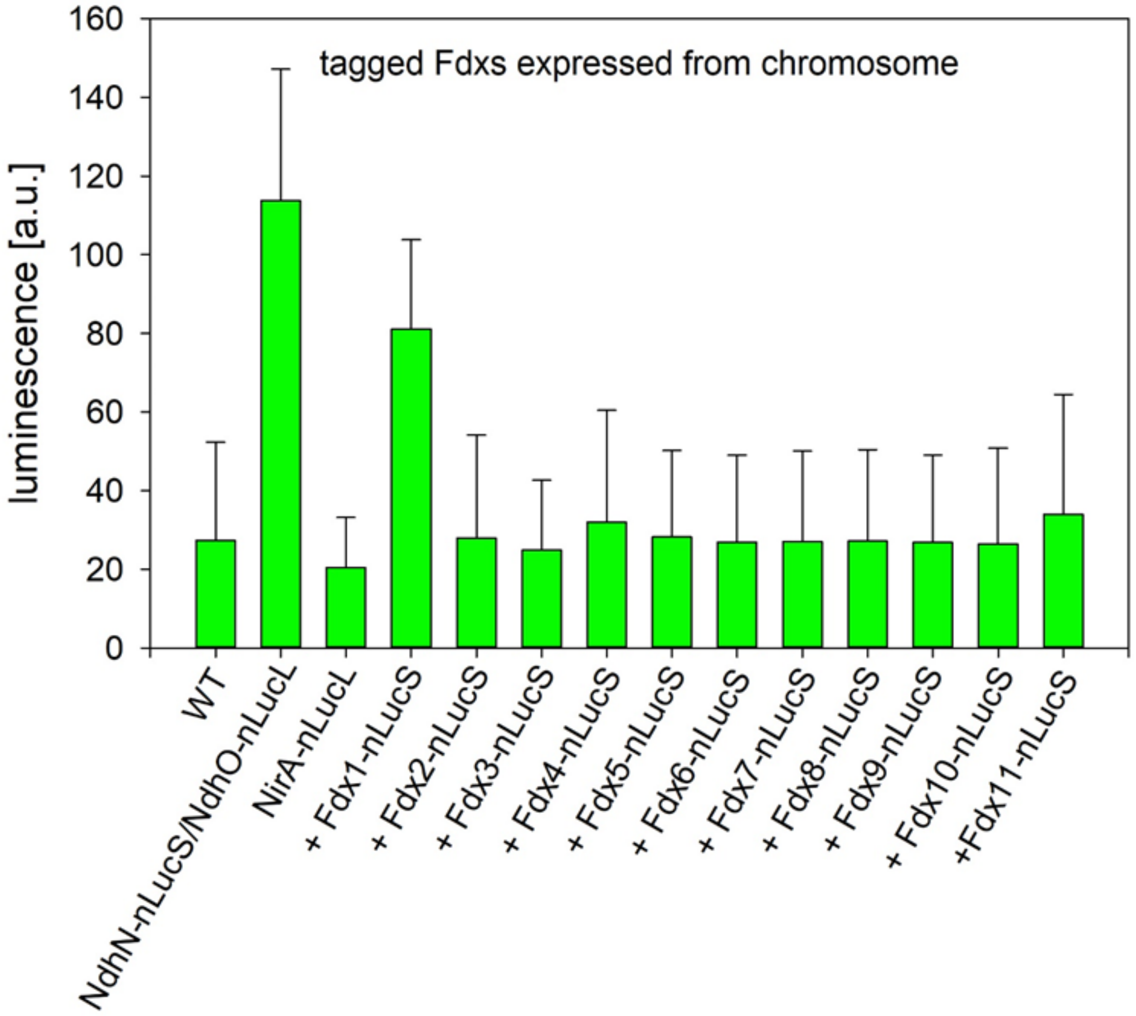


**Fig. S13:** *In vivo* interaction of nitrite reductase with Fdxs1-11 expressed under the control of their native promoters. In all strains the nitrite reductase and Fdxs were tagged with the LgBit subunit (nLucL) or the SmBit subunit (nLucS), respectively, at the genome level (integration via homologous recombination). The NdhO-nLucL / NdhS-nLucS expressing strain was measured as a positive control. Plots are the means and standard deviations of three independent experiments.

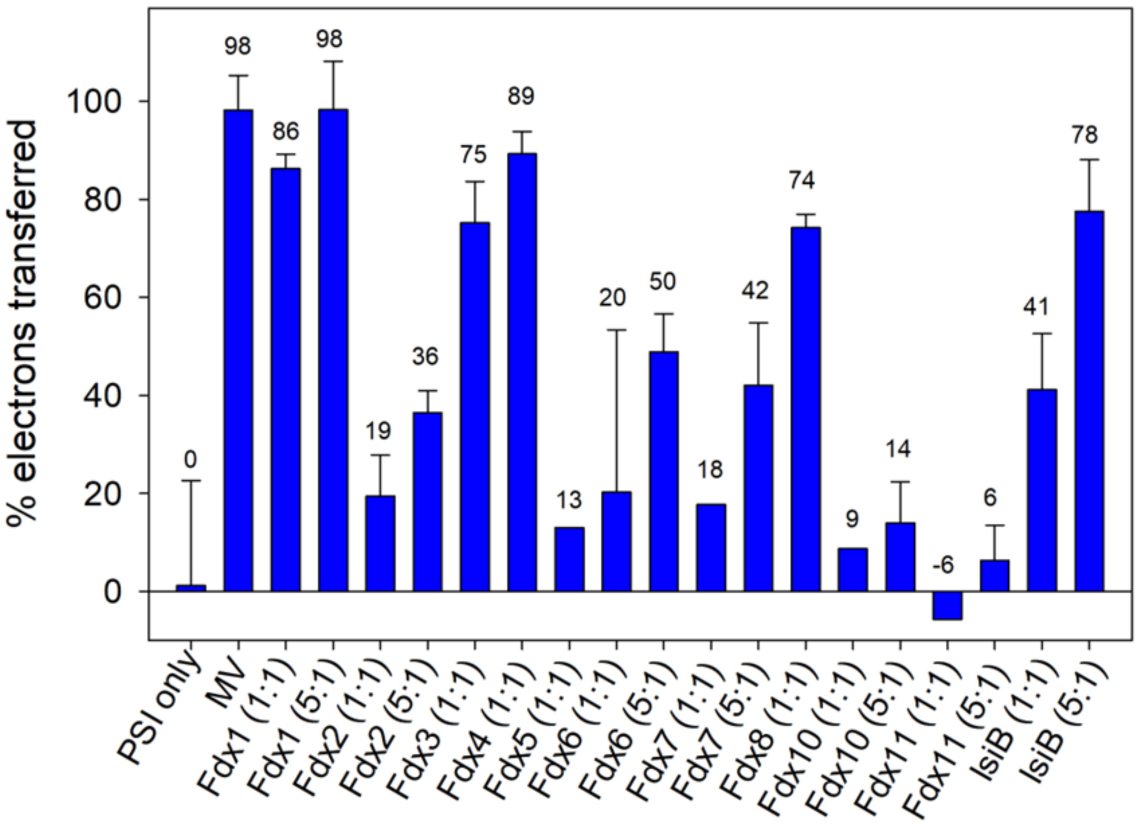


**Fig. S14:** Electron transfer from photosystem I (PSI) to purified Fdxs. Electron transfer was assessed by $P700^+$ reduction kinetics measurements with PSI at a concentration of 500 nM and Fdx in equal concentration (500 nM, 1:1) or in excess (2500 nM, 5:1). The method for PSI purification is described in **Supporting Text 4**. PSI only and PSI with added methyl viologen (1 mM) were measured as negative and positive controls, respectively. Fdx9-Strep (Fig. S6) was measured, however, due to the insufficient quality of this sample and its oxygen sensitivity, we denote "n.d." in Table 2. A value of above 30% of electrons transferred was chosen as the cutoff and resulted in a "+" denotation in Table 2, when this value was observed for both ratios. When this threshold was only surpassed for the 5:1 ratio, this resulted in a "+/-" denotation for reactivity with PSI in Table 2.

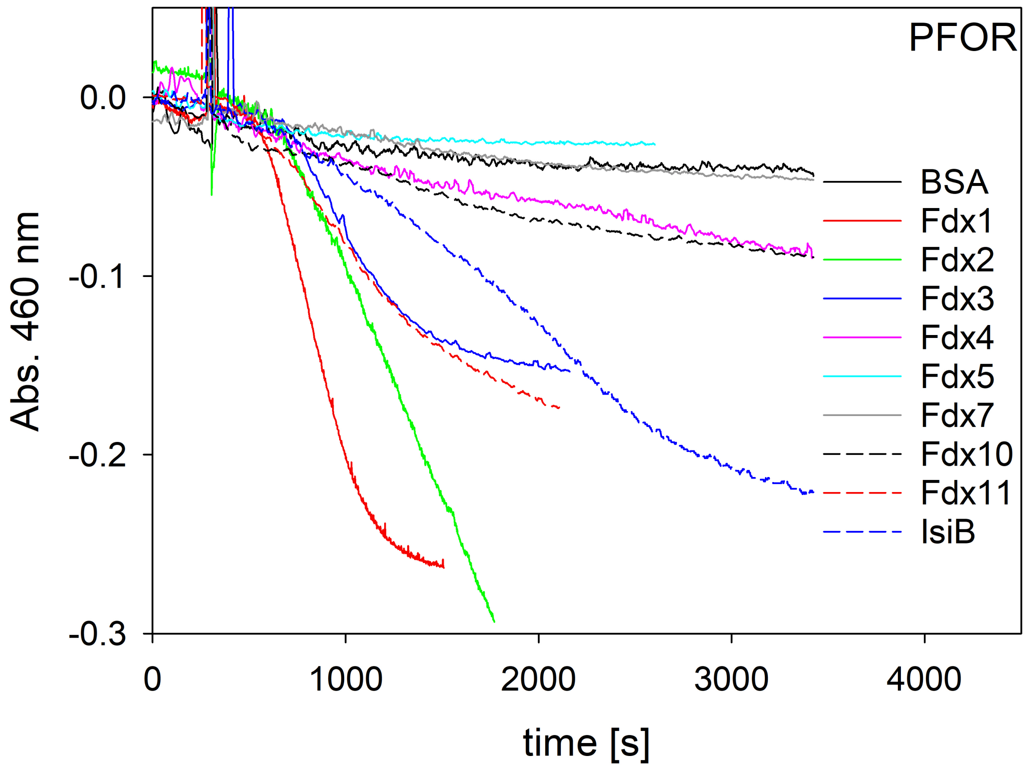


**Fig. S15:** Electron transfer from pyruvate ferredoxin oxidoreductase (PFOR) to purified Fdxs. To assess Fdx reduction, Fdx absorbance was monitored at 460 nm in the PFOR/Fdx reduction assay. Fdx5 could not be determined, because there is no detectable change in absorption for this protein (Fig. S7). Fdx6 and Fdx8 could not be tested due to insufficient material. In these measurements the Fdx9* sample (Fig. S6) was measured. However, due to the insufficient quality of this particular sample and its oxygen sensitivity, we denote “n.d.” in Table 2.

GACTACTATAGGGCGAATTGGGTACCATATGAAGAAGCGTGTGACCCTGACTTTTCCCC
GTAGTGCGGTGCAAATGCCCGTCACTTATCGGCTGGCCAAGGATTTTAATATTGCCGCC
AATATTATCCGAGCCCAGGTGGCCCCCAATCAGGTAGGCAAAGTAGTACTGGAATTGTC
CGGGGACATTGACCAACTGGAAGCGTCTTTGGAGTGGATGCGCTCCCAGAGTATTGAAG
TTTCCTTGGCTAGTCGGGAAATTGTCATTGATGACCAAAGTTGTGTAGATTGTGGCCTC
TGTACCGGGGTTTGCCCTACGGAGGCTCTGAGTTTAGATCCGGATAGTTTTCGCTTGAT
GTTCCGTCGTTCCCGTTGTGTGGTCTGTGAGCAATGTATTCCTTCCTGCCCCGTCCAGG
CGATCGCCACTAATTTCCCCATCGGAGGATCCGAAAACCTGTATTTTCAGGGCGGCGCT
AGCCCCGGGCATCACCATCACCATCACTAAGATATCTAGGAATTCAGCTCCAGCTTTTG
TTCCCTTTAGT

**Fig. S16.** Fragment ordered from Genscript for the generation of the pMB0122 vector. The following features are indicated: green highlight, sequence overlapping with the pBluescript SK+ vector; yellow highlight, coding sequence of *Synechocystis* Fdx9; blue highlight, a linker sequence containing a *Bam*HI restriction site and TEV cleavage site; cyan highlight, NheI restriction site; gray highlight, SmaI restriction site; underlined yellow highlight, 6-His-tag coding sequence; purple highlight, TAA and TAG stop codons; red highlight, EcoRV restriction site; black highlight, EcoRI restriction site.

Fdx6 (ssl2559; codon-optimized):
CATATGAATAATTGCGTAATTTCTTTTCCTCAAACTAAATTTTTGCCCCTATCACTAGAATTTAACGCTTG
TCTAGCGGAATATTTAACCCCAGACAATTCGCCAATTTTATTCGGTTGTCGCACAGGATTATGTGGTACCT
GTCTAGTGAAAGTCGTGGGGGAAATCCTCTCCCCAGAGGCCGAGGAAAGGGAAATTTTGGCTATTCTGGCT
CCTGATGATGTTCAGGCTAGATTGGCTTGTCAAATTAAGTTAACAGGAGATATTGCCATTAGAGCTTATCA
AAGTGATGAAATTCCCATCGGAGGATCC

Fdx8 (ssr3184; codon-optimized):
CATATGCCCCATACCATTGTTACGGAAACCTGCGAAGGCGTGGCCGACTGCGTTGAAGCCTGCCCCGTAGC
CTGCATTCACCCCGGAGACGGGAAAAACACCATTGGCACCGATTGGTACTGGATTGACTTTGCCACCTGCA
TCGATTGCGGCATCTGCCTCCAAGTTTGCCCGGTGGAAGGGGCCATTCTGCCGGAGGAACGGCCTGATTTA
CAGAAAAGCCCTGCCCCCATCGGAGGATCC

Fdx9 (slr2059; codon-optimized):
CATATGAAGAAGCGTGTGACCCTGACTTTTCCCCGTAGTGCGGTGCAAATGCCCGTCACTTATCGGCTGGC
CAAGGATTTTAATATTGCCGCCAATATTATCCGAGCCCAGGTGGCCCCCAATCAGGTAGGCAAAGTAGTAC
TGGAATTGTCCGGGGACATTGACCAACTGGAAGCGTCTTTGGAGTGGATGCGCTCCCAGAGTATTGAAGTT
TCCTTGGCTAGTCGGGAAATTGTCATTGATGACCAAAGTTGTGTAGATTGTGGCCTCTGTACCGGGGTTTG
CCCTACGGAGGCTCTGAGTTTAGATCCGGATAGTTTTCGCTTGATGTTCCGTCGTTCCCGTTGTGTGGTCT
GTGAGCAATGTATTCCTTCCTGCCCCGTCCAGGCGATCGCCACTAATTTCCCCATCGGAGGATCC

Fdx10 (sll1584; codon-optimized):
CATATGGAAAACCCTGTCCCCATAACTCCCTCAGAAACCATTGTCGATAGTTGCCAACGTTTAGGCCTTGG
GCGCATTCAACGGCATCTGTTTCTCTGTTGTGACCAAACAAAACCGAAATGCTGTAGCAAAGAAGATAGCC
TCGCCACCTGGGATTATCTGAAAAAGCGTTTGCCGGAGTTGGGGCTGGATTGCACTCAGTCCAGTCGTGAT
GGGAACATTTTCCGCACTAAGGCTAATTGTTTGCGGGTTTGCCAGCAGGGGCCAATTTTACTGGTATATCC
AGAGGGCATTTGGTACCGGAACGTGACCCCCACAGTAATGGAAAAAATTCTCCAAGAGCATATTTTGCAAA
ACCGTCCGGTGGAGGAGTATCGCTTTTTTACCCACCCTCTTTCGCATTTACCCATCGGAGGATCC

**Fig. S17.** Codon-optimized sequences of various Fdxs (Genscript). Added restriction sites are indicated: NdeI, *Bam*HI. Additional linker sequence is underlined.

### Supporting Tables

**Table S1.** A comparison of the surface charges on all *Synechocystis* Fdxs studied in this work. The surface charge predictions were obtained in Pymol using the APBS surface charge calculation plugin.

| Protein | Total surface residues | Positively charged aa | Negatively charged aa | Total charge @ pH 7 | List of charged surface residues |
|---|---|---|---|---|---|
| Fdx1 | 80 | 5 | 20 | -15.88 | KDEEDDDEEDRKDDDDDEDEHKEED |
| Fdx2 | 103 | 14 | 21 | -9.68 | RHRDREKDDDRHEDERRHEDRDEEDEDEERRRDED |
| Fdx3 | 91 | 6 | 16 | -10.89 | EHEDKDDEEEDEERDKEKEDER |
| Fdx4 | 85 | 7 | 17 | -11.79 | DEEDEEEDRREEHDKDKEEDKHEE |
| Fdx5 | 141 | 21 | 25 | -5.78 | KKDDKEDDRKERHERRREHRRRDEEEEDEERREERDEKKRDDKDDE |
| Fdx6 | 85 | 7 | 12 | -4.99 | KEEDRKEEEREDDRKDRDE |
| Fdx7 | 82 | 14 | 14 | -5.39 | ERDKHEEHRDDEEDDHDEKHEEKRKHRH |
| Fdx8 | 70 | 5 | 12 | -8.79 | HEEDEHDKDDDEEERDK |
| Fdx9 | 118 | 14 | 15 | -0.99 | KKRRRKDRKEDDEEREREDDDEDDRRRRE |
| Fdx10 | 114 | 20 | 15 | +2.31 | EEDRRRHDKKKEDDKKREDRDRKREREKEREERHH |
| Fdx11 | 68 | 2 | 12 | -9.99 | DDREDEREEDEEDD |
| Fdx12 | 81 | 13 | 8 | +5 | KEKDREKDKKEEKDRRKKDKK |

**Table S2.** Summary for Fdx1 to Fdx12 and flavodoxin (IsiB) purifications. Fdx8 and Fdx9 were prepared using two different expression plasmids and strains (ΔiscR BL21 strain is described in (17)). Fdx8-Strep was not analyzed by SDS-PAGE (Fig. S6). The purification method for Fdx9-Strep performed as described in (18). Fdx6, Fdx9 and Fdx10 were purified from combinations of separate expressions. Fdx12 could not be expressed as soluble protein and was therefore not purified in this study. Protein concentrations were determined by Bradford assay, and the relative abundance was determined by LC/MS (data not shown; for the method see **Supporting Text 2**). Buffer A: 25 mM NaPO4, pH=7.0; 50 mM NaCl; 5% (v/v) glycerol; Buffer B: 50 mM Tris pH 8.5, 50 mM NaCl, 5% glycerol. Abbreviations: Fed6co, Fdx6 codon optimized sequence; TEV, TEV protease recognition site; GST, glutathione-S-transferase affinity-tag; 6xHis, 6x histidine affinity-tag; MBP, maltose-binding protein; Strep, Strep affinity-tag; n.d., not determined.

| **Protein** | **Expression Plasmid** | ***E. coli* Expression strain** | **Protein [µM]** | **relative abundance** | **adjusted protein [µM]** | **buffer** |
|---|---|---|---|---|---|---|
| Fdx1 | Fed1-TEV-GST-6xHis | KRX | 652 | 8.53% | 56 | A |
| Fdx2 | Fed2-TEV-GST-6xHis | KRX | 1326 | 90.64% | 1202 | A |
| Fdx3 | Fed3-TEV-GST-6xHis | KRX | 1086 | 69.57% | 755 | A |
| Fdx4 | Fed4-TEV-GST-6xHis | KRX | 1382 | 97.38% | 1346 | A |
| Fdx5 | Fed5-TEV-GST-6xHis | KRX | 1419 | 99.95% | 1418 | A |
| Fdx6 | Fed6-TEV-GST-6xHis<br>Fed6co-TEV-MBP-His | KRX<br>C41 (DE3) | 270 | 74.08% | 200 | A |
| Fdx7 | Fed7-TEV-GST-6xHis | KRX | 934 | 98.37% | 919 | A |
| Fdx8 | Fed8co-TEV-MBP-His | KRX | 955 | n.d. | n.d. | B |
| Fdx8* | Fed8-Strep | ΔiscR BL21 | n.d. | n.d. | n.d. | B |
| Fdx9 | Fed9-TEV-GST-6xHis<br>Fed9co-TEV-MBP-His | KRX<br>KRX | 247 | n.d. | n.d. | A |
| Fdx9* | Fed9-Strep | BL21 (DE3) | 116 | 84.93% | 99 | B |
| Fdx10 | Fed10-TEV-GST-6xHis<br>Fed10co-TEV-MBP-His | KRX<br>KRX | 518 | 98.16% | 508 | A |
| Fdx11 | Fed11-TEV-GST-6xHis | KRX | 1603 | 79.40% | 1273 | A |
| Fdx12 | Fed12-TEV-GST-6xHis | No soluble protein expression in *E. coli* | | | | |
| IsiB | IsiB-TEV-GST-6xHis | Rosetta 2<br>pLysS (DE3) | 771 | 92.24% | 711 | A |

**Table S3.** EPR properties of the reduced *Synechocystis* Fdx signals presented in Fig. 3.

| Protein | *g*-values | Rhombicity (%)[a] | Fe-S cluster type | Reference |
|---|---|---|---|---|
| Fdx1 | $g$ = 2.048, 1.955, 1.877; $g_{av}$ = 1.96 | 89 | [2Fe-2S] | this work, (19) |
| Fdx2 | $g$ = 2.048, 1.957, 1.869; $g_{av}$ = 1.96 | 98 | [2Fe-2S] | this work |
| Fdx3 | $g$ = 2.044, 1.952, 1.868; $g_{av}$ = 1.95 | 94 | [2Fe-2S] | this work |
| Fdx4[b] | $g$ = 2.054, 1.963, 1.878; $g_{av}$ = 1.97 | 96 | [2Fe-2S] | this work, (20) |
| Fdx5 | n.d. | n.d. | n.d. | (20) |
| Fdx6 | $g$ = 2.040, 1.946, 1.883; $g_{av}$ = 1.96 | 75 | [2Fe-2S] | this work |
| Fdx7[c] | $g$ = 2.068, 1.926, 1.888; $g_{av}$ = 1.96<br>$g$ = 2.058, 1.932, 1.869; $g_{av}$ = 1.95 | 35<br>60 | [4Fe-4S] | this work |
| Fdx8[d] | $g$ = 2.068, 1.932, 1.871; $g_{av}$ = 1.96<br>$g$ = 2.058, 1.929, 1.890; $g_{av}$ = 1.96 | 55<br>39 | 2x[4Fe-4S] | this work |
| Fdx9[e] | $g$ = 2.07, 2.05, 2.01, 1.99, 1.94 | u.d. | 2x[4Fe-4S] | this work, (18) |
| Fdx10 | $g$ = 2.002, 1.948, 1.919; $g_{av}$ = 1.96 | 64 | [2Fe-2S] | this work |
| Fdx11 | $g$ = 2.016, 1.941, 1.930; $g_{av}$ = 1.96 | 20 | [2Fe-2S] | this work |

[a]Calculated according to the following equation: percent rhombicity = $300^*((g_y\text{-}g_x)/(2g_z\text{-}g_y\text{-}g_x))$, where $g_z > g_y > g_x$ (21). [b]Fdx4 is homologous to Fdx2 from *Thermosynechococcus elongatus* and displayed a similar EPR signal at $g$ = 2.051, 1.964, 1.869 (20). No EPR signal could be obtained for Fdx5. [c]The spectrum of Fdx7 was simulated with two overlapping rhombic signals, with the total simulation consisting of 64% contribution of the $g$ = 2.068 signal and 36% contribution of the $g$ = 2.058 signal, along with a very minor contribution of a species at $g$ = 2.003 (<1%). [d]The spectrum of Fdx8 was simulated with 2 overlapping rhombic signals, with the total simulation consisting of 68% contribution of the $g$ = 2.068 signal and 32% contribution of the $g$ = 2.058 signal, along with a very minor contribution of a species at $g$ = 2.003 (<1%). [e]The $g$-values listed here for Fdx9 give the general positions of spectral features and were not obtained from simulation; initial attempts at simulating the complex spectrum of Fdx9 with and without the addition of spin-spin coupling are reported in (18). Abbreviations: $g_{av}$, $g$-value average, n.d., not detected; u.d., undetermined; n.a., not available.

**Table S4.** Primers used in this work.

| # | primer name | internal reference | sequence | used for |
|---|---|---|---|---|
| 1 | Syn_ssl0020_Fed1-fw | MB01_B4 | 5'-GCGCATATGGCATCCTATACCGTTAAATTGATCACC-3' | cloning of expression construct |
| 2 | Syn_ssl0020_Fed1-rev | MB01_C8 | 5'-GCGGGATCCTCCGATGGGGTAGAGGTCTTCTTCTTTGTGGGTTTCAAT-3' | cloning of expression construct |
| 3 | Syn_sll1382_Fed2-fw | MB01_B5 | 5'-GCGCATATGTCCCGTTCCCACCGAGTTCT-3' | cloning of expression construct |
| 4 | Syn_sll1382_Fed2-rev | MB01_C9 | 5'-GCGGGATCCTCCGATGGGGTCCTCATCTAAAGGCAAACCTAATCG-3' | cloning of expression construct |
| 5 | Syn_slr1828_Fed3-fw | MB01_B6 | 5'-GCGCATATGGTTAACACCTACACCGCCGAAAT-3' | cloning of expression construct |
| 6 | Syn_slr1828_Fed3-rev | MB01_D1 | 5'-GCGGGATCCTCCGATGGGACCTTGGCCGCCGAACTGCC-3' | cloning of expression construct |
| 7 | Syn_slr0150_Fed4-fw | MB01_B7 | 5'-GCGCATATGGGTGCAATTTATTCCGTCAATTTAGTCAAT-3' | cloning of expression construct |
| 8 | Syn_slr0150_Fed4-rev | MB01_D2 | 5'-GCGGGATCCTCCGATGGGACCAAACAACGCTTCCTCTTGGTGAG-3' | cloning of expression construct |
| 9 | Syn_slr0148_Fed5-fw | MB01_B8 | 5'-GCGCATATGGCTAAAACTATTAAGCTCGACCCCAT-3' | cloning of expression construct |
| 10 | Syn_slr0148_Fed5-rev | MB01_D3 | 5'-GCGGGATCCTCCGATGGGTTCATCGGTATCGTTGACAATCTGAATTTTGG-3' | cloning of expression construct |
| 11 | Syn_ssl2559_Fed6-fw | MB01_B9 | 5'-GCGCATATGAATAATTGCGTAATTTCTTTTCCTCAAACTAAATTTTTG-3' | cloning of expression construct |
| 12 | Syn_ssl2559_Fed6-rev | MB01_D4 | 5'-GCGGGATCCTCCGATGGGAATTTCATCACTTTGATAAGCTCTAATGGCAATATCT-3' | cloning of expression construct |
| 13 | Syn_sll0662_Fed7-fw | MB01_C1 | 5'-GCGCATATGGTGATAGCTGATCTAAATTTTCCTCCC-3' | cloning of expression construct |
| 14 | Syn_sll0662_Fed7-rev | MB01_D5 | 5'-GCGGGATCCTCCGATGGGCAAATGGGGATTGATTTGCGGGTAACC-3' | cloning of expression construct |
| 15 | Syn_ssr3184_Fed8-fw | MB01_C2 | 5'-GCGCATATGCCCCATACCATTGTTACGGAAACC-3' | cloning of expression construct |
| 16 | Syn_ssr3184_Fed8-rev | MB01_D6 | 5'-GCGGGATCCTCCGATGGGGGCAGGGCTTTTCTGTAAATCAGGC-3' | cloning of expression construct |
| 17 | Syn_slr2059_Fed9-fw | MB01_C3 | 5'-GCGCATATGAAGAAGCGTGTGACCCTGACTTTT-3' | cloning of expression construct |
| 18 | Syn_slr2059_Fed9-rev | MB01_D7 | 5'-GCGGGATCCTCCGATGGGGAAATTAGTGGCGATCGCCTGGAC-3' | cloning of expression construct |
| 19 | Syn_sll1584_Fed10-fw | MB01_C4 | 5'-GCGCATATGGAAAACCCTGTCCCCATAACTCC-3' | cloning of expression construct |
| 20 | Syn_sll1584_Fed10-rev | MB01_D8 | 5'-GCGGGATCCTCCGATGGGTAAATGCGAAAGAGGGTGGGTAAAAAAGC-3' | cloning of expression construct |
| 21 | Syn_ssl3044_Fed11-fw | MB01_C5 | 5'-GCGCATATGACCATTACCTTTGTTAAAGAGCAGAAGG-3' | cloning of expression construct |
| 22 | Syn_ssl3044_Fed11-rev | MB01_D9 | 5'-GCGGGATCCTCCGATGGGGCCTTTGGGTTTGGTGTTCACACTCA-3' | cloning of expression construct |
| 23 | Syn_ssr1041_Fed12-fw | MB10_A2 | 5'-TTTAACTTTAAGAAGGAGATATACATATGATGGCAGTAACGATACACTTTTTGC-3' | cloning of expression construct |
| 24 | Syn_ssr1041_Fed12-rev | MB10_A3 | 5'-CCTGAAAATACAGGTTTTCGGATCCTCCGATGGGCCAGGTGAGATCATCATAAAGATTAATTTCTAATTCCTG-3' | cloning of expression construct |
| 25 | Syn_sll0248_IsiB-fw | MB02_A3 | 5'-GTTTAACTTTAAGAAGGAGATATACATATGACAAAAATTGGACTTTTTTACGGTAC-3' | cloning of expression construct |
| 26 | Syn_sll0248_IsiB-rev | MB02_A4 | 5'-GCCGCCCTGAAAATACAGGTTTTCGGATCCTCCGATGGGGGATTGCAAAATTGGTTTAATTTCAC-3' | cloning of expression construct |
| 27 | MBP-fw | Common | 5'-GCTAGCATGAAAATCGAAGAAGGTAAACTGGTAATCTG-3' | cloning of expression construct |
| 28 | MBP-rev | Common | 5'-CCCGGGAGTCTGCGCGTCTTTCAGGGCTT-3' | cloning of expression construct |
| 29 | Fed1-His_1-fw | MB03_F1 | 5'-GACTACTATAGGGCGAATTGGGTACCTCGAGATGGCATCCTATACCGTTAAATTGATCACC-3' | generation of C-entry plasmid, mutant confirmation |
| 30 | Fed1-His_2-rev | MB03_F2 | 5'-GAATTCCTCGAGGGATCCTCCGATGGGGTAGAGGTCTTCTTCTTTGTGGGTTTCA-3' | generation of C-entry plasmid |

| # | primer name | internal reference | sequence | used for |
|---|---|---|---|---|
| 31 | Fed1-His_3-fw | MB03_F3 | 5'-CCCATCGGAGGATCCCTCGAGGAATTCGGGTAATAATGCTGGCCATGGGCTA-3' | generation of C-entry plasmid |
| 32 | Fed1-His_4-rev | MB03_F4 | 5'-ACTAAAGGGAACAAAAGCTGGAGCTCTGGAAATTGCTTTACTAAATGTGCCCC-3' | generation of C-entry plasmid, mutant confirmation |
| 33 | Fed2-His_1-fw | MB03_F5 | 5'-GACTACTATAGGGCGAATTGGGTACCTCGAGATGTCCCGTTCCCACCGAGTTCT-3' | generation of C-entry plasmid, mutant confirmation |
| 34 | Fed2-His_2-rev | MB03_F6 | 5'-GAATTCCTCGAGGGATCCTCCGATGGGGTCCTCATCTAAAGGCAAACCTAATCG-3' | generation of C-entry plasmid |
| 35 | Fed2-His_3-fw | MB03_F7 | 5'-CCCATCGGAGGATCCCTCGAGGAATTCGTAGGCTACAACTACAACCTGATTTTCC-3' | generation of C-entry plasmid |
| 36 | Fed2-His_4-rev | MB03_F8 | 5'-ACTAAAGGGAACAAAAGCTGGAGCTGGGAATGTCTTGGATCAGTAATCCAATG-3' | generation of C-entry plasmid, mutant confirmation |
| 37 | Fed3-His_1-fw | MB03_F9 | 5'-GACTACTATAGGGCGAATTGGGTACCTCGAGATGGTTAACACCTACACCGCCGAAATC-3' | generation of C-entry plasmid, mutant confirmation |
| 38 | Fed3-His_2-rev | MB03_G1 | 5'-GAATTCCTCGAGGGATCCTCCGATGGGACCTTGGCCGCCGAACTGCC-3' | generation of C-entry plasmid |
| 39 | Fed3-His_3-fw | MB03_G2 | 5'-CCCATCGGAGGATCCCTCGAGGAATTCATTCGGCTGGAATTCTCCCTTCCC-3' | generation of C-entry plasmid |
| 40 | Fed3-His_4-rev | MB03_G3 | 5'-ACTAAAGGGAACAAAAGCTGGAGCTGCTGTTTTAGGTAGCCCTGCACTG-3' | generation of C-entry plasmid, mutant confirmation |
| 41 | Fed4-His_1-fw | MB03_G4 | 5'-GACTACTATAGGGCGAATTGGGTACCTCGAGATGGGTGCAATTTATTCCGTCAATTTAGTCA-3' | generation of C-entry plasmid, mutant confirmation |
| 42 | Fed4-His_2-rev | MB03_G5 | 5'-GAATTCCTCGAGGGATCCTCCGATGGGACCAAACAACGCTTCCTCTTGGTGAGT-3' | generation of C-entry plasmid |
| 43 | Fed4-His_3-fw | MB03_G6 | 5'-CCCATCGGAGGATCCCTCGAGGAATTCTCAGCCACCGGCTACAACTCCG-3' | generation of C-entry plasmid |
| 44 | Fed4-His_4-rev | MB03_G7 | 5'-ACTAAAGGGAACAAAAGCTGGAGCTGCAGCGGCTTGGAAAGCTTCCAAC-3' | generation of C-entry plasmid, mutant confirmation |
| 45 | Fed5-His_1-fw | MB03_G8 | 5'-GACTACTATAGGGCGAATTGGGTACCTCGAGGTGGCTAAAACTATTAAGCTCGACCC-3' | generation of C-entry plasmid, mutant confirmation |
| 46 | Fed5-His_2-rev | MB03_G9 | 5'-GAATTCCTCGAGGGATCCTCCGATGGGTTCATCGGTATCGTTGACAATCTGAATTTTG-3' | generation of C-entry plasmid |
| 47 | Fed5-His_3-fw | MB03_H1 | 5'-CCCATCGGAGGATCCCTCGAGGAATTCTTTCCTGGCTGACTGTGGCGGTT-3' | generation of C-entry plasmid |
| 48 | Fed5-His_4-rev | MB03_H2 | 5'-ACTAAAGGGAACAAAAGCTGGAGCTAATAGACCCTCCCGCAGACGGT-3' | generation of C-entry plasmid, mutant confirmation |
| 49 | Fed6-His_1-fw | MB03_H3 | 5'-GACTACTATAGGGCGAATTGGGTACCTCGAGGCGTAATTTCTTTTCCTCAAACTAAATTTTTGCC-3' | generation of C-entry plasmid, mutant confirmation |
| 50 | Fed6-His_2-rev | MB03_H4 | 5'-GAATTCCTCGAGGGATCCTCCGATGGGAATTTCATCACTTTGATAAGCTCTAATGGCAATATC-3' | generation of C-entry plasmid |
| 51 | Fed6-His_3-fw | MB03_H5 | 5'-CCCATCGGAGGATCCCTCGAGGAATTCAGATTTTTGGTATGTGGTGGCTGAAAGTAG-3' | generation of C-entry plasmid |
| 52 | Fed6-His_4-rev | MB03_H6 | 5'-ACTAAAGGGAACAAAAGCTGGAGCTGCAGTTGGTGACATTATTAGCAAATCGATG-3' | generation of C-entry plasmid, mutant confirmation |

| # | primer name | internal reference | sequence | used for |
|---|---|---|---|---|
| 53 | Fed7-His_1-fw | MB03_H7 | 5'-GACTACTATAGGGCGAATTGGGTACCTCGAGATGGTGATAGCTGATCTAAATTTTCCTCCC-3' | generation of C-entry plasmid, mutant confirmation |
| 54 | Fed7-His_2-rev | MB03_H8 | 5'-GAATTCCTCGAGGGATCCTCCGATGGGCAAATGGGGATTGATTTGCGGGTAAC-3' | generation of C-entry plasmid |
| 55 | Fed7-His_3-fw | MB03_H9 | 5'-CCCATCGGAGGATCCCTCGAGGAATTCTTAATTGGGTGATGGAATCTAATCGAAATCAG-3' | generation of C-entry plasmid |
| 56 | Fed7-His_4-rev | MB03_I1 | 5'-ACTAAAGGGAACAAAAGCTGGAGCTGGTATTTGAAACGGTGGTGTTATTTTTGCTG-3' | generation of C-entry plasmid, mutant confirmation |
| 57 | Fed8-His_1-fw | MB03_I2 | 5'-GACTACTATAGGGCGAATTGGGTACCTCGAGGTTGAACGTCCGCTACCCTTAGAAC-3' | generation of C-entry plasmid, mutant confirmation |
| 58 | Fed8-His_2-rev | MB03_I3 | 5'-GAATTCCTCGAGGGATCCTCCGATGGGGGCAGGGCTTTTCTGTAAATCAGGC-3' | generation of C-entry plasmid |
| 59 | Fed8-His_3-fw | MB03_I4 | 5'-CCCATCGGAGGATCCCTCGAGGAATTCGGTTGGGAGGGGTCTAACTGC-3' | generation of C-entry plasmid |
| 60 | Fed8-His_4-rev | MB03_I5 | 5'-ACTAAAGGGAACAAAAGCTGGAGCTGGTAGGACCCTGGTGTAGAAAGG-3' | generation of C-entry plasmid, mutant confirmation |
| 61 | Fed9-His_1-fw | MB03_I6 | 5'-GACTACTATAGGGCGAATTGGGTACCTCGAGATGAAGAAGCGTGTGACCCTGACTTTT-3' | generation of C-entry plasmid, mutant confirmation |
| 62 | Fed9-His_2-rev | MB03_I7 | 5'-GAATTCCTCGAGGGATCCTCCGATGGGGAAATTAGTGGCGATCGCCTGGAC-3' | generation of C-entry plasmid |
| 63 | Fed9-His_3-fw | MB03_I8 | 5'-CCCATCGGAGGATCCCTCGAGGAATTCGTTCTCAAACGGCGATCGCCAAAATTG-3' | generation of C-entry plasmid |
| 64 | Fed9-His_4-rev | MB03_I9 | 5'-ACTAAAGGGAACAAAAGCTGGAGCTGTAAGCCAGGGCGGTCTAATTTCCC-3' | generation of C-entry plasmid, mutant confirmation |
| 65 | Fed10-His_1-fw | MB04_A1 | 5'-GACTACTATAGGGCGAATTGGGTACCTCGAGATGGAAAACCCTGTCCCCATAACTCC-3' | generation of C-entry plasmid, mutant confirmation |
| 66 | Fed10-His_2-rev | MB04_A2 | 5'-GAATTCCTCGAGGGATCCTCCGATGGGTAAATGCGAAAGAGGGTGGGTAAAAAAGC-3' | generation of C-entry plasmid |
| 67 | Fed10-His_3-fw | MB04_A3 | 5'-CCCATCGGAGGATCCCTCGAGGAATTCATTATCAAAACCGCTATAGAAACGAGTCGG-3' | generation of C-entry plasmid |
| 68 | Fed10-His_4-rev | MB04_A4 | 5'-ACTAAAGGGAACAAAAGCTGGAGCTGAGGAAACCAACCTTTTAATCGCCGAAAA-3' | generation of C-entry plasmid, mutant confirmation |
| 69 | Fed11-His_1-fw | MB04_A5 | 5'-GACTACTATAGGGCGAATTGGGTACGTGACCATTACCTTTGTTAAAGAGCAGAAG-3' | generation of C-entry plasmid, mutant confirmation |
| 70 | Fed11-His_2-rev | MB04_A6 | 5'-GAATTCCTCGAGGGATCCTCCGATGGGGCCTTTGGGTTTGGTGTTCACACTCA-3' | generation of C-entry plasmid |
| 71 | Fed11-His_3-fw | MB04_A7 | 5'-CCCATCGGAGGATCCCTCGAGGAATTCGGGCGATCACCGGAGGCCA-3' | generation of C-entry plasmid |
| 72 | Fed11-His_4-rev | MB04_A8 | 5'-ACTAAAGGGAACAAAAGCTGGAGCTGTTGTACTTTATAGTGCTGTAGTATCCCG-3' | generation of C-entry plasmid, mutant confirmation |
| 73 | Gent-fw | Common | 5'-GGTTCGTGCCTTCATCCGTCGAC-3' | generation of toolkit vector |
| 74 | Gent-rev | Common | 5'-CGCCACCTAACAATTCGGTCGAC-3' | generation of toolkit vector |
| 75 | NirA-fw | MB05_H7 | 5'-GACGGTGCCTAAGCCCGTGCGGATCC-3' | mutant confirmation |
| 76 | NirA-rev | MB05_H6 | 5'-GTAAGTTACCTACCTATCCACGAACCAT-3' | mutant confirmation |

| # | primer name | internal reference | Sequence | used for |
|---|---|---|---|---|
| 77 | Fed1-nLuc-pSSR-fw | MB09_F2 | 5'-GAGTAGTGGAGGTTACCATATGGCATCCTATACCGTTAAATTGATCAC-3' | rhamnose inducible expression |
| 78 | Fed1-nLuc-pSSR-rev | MB09_F3 | 5'-CGGCAGCAGATCTCATGTAGAGGTCTTCTTCTTTGTGGGT-3' | rhamnose inducible expression |
| 79 | Fed2-nLuc-pSSR-fw | MB09_F4 | 5'-GAGTAGTGGAGGTTACCATATGTCCCGTTCCCACCGAGTTCT-3' | rhamnose inducible expression |
| 80 | Fed2-nLuc-pSSR-rev | MB09_F5 | 5'-CGGCAGCAGATCTCATGTCCTCATCTAAAGGCAAACCTAAT-3' | rhamnose inducible expression |
| 81 | Fed3-nLuc-pSSR-fw | MB09_F6 | 5'-GAGTAGTGGAGGTTACCATATGGTTAACACCTACACCGCC-3' | rhamnose inducible expression |
| 82 | Fed3-nLuc-pSSR-rev | MB09_F7 | 5'-CGGCAGCAGATCTCATACCTTGGCCGCCGAA-3' | rhamnose inducible expression |
| 83 | Fed4-nLuc-pSSR-fw | MB09_F8 | 5'-GAGTAGTGGAGGTTACCATATGGGTGCAATTTATTCCGTCAATTTAGT-3' | rhamnose inducible expression |
| 84 | Fed4-nLuc-pSSR-rev | MB09_F9 | 5'-CGGCAGCAGATCTCATACCAAACAACGCTTCCTCTTG-3' | rhamnose inducible expression |
| 85 | Fed5-nLuc-pSSR-fw | MB09_G1 | 5'-GAGTAGTGGAGGTTACCATATGGCTAAAACTATTAAGCTCGACCC-3' | rhamnose inducible expression |
| 86 | Fed5-nLuc-pSSR-rev | MB09_G2 | 5'-CGGCAGCAGATCTCATTTCATCGGTATCGTTGACAATCTGAATTTTG-3' | rhamnose inducible expression |
| 87 | Fed6-nLuc-pSSR-fw | MB09_G3 | 5'-GAGTAGTGGAGGTTACCATATGAATAATTGCGTAATTTCTTTTCCTCAAACTAAATTT-3' | rhamnose inducible expression |
| 88 | Fed6-nLuc-pSSR-rev | MB09_G4 | 5'-CGGCAGCAGATCTCATAATTTCATCACTTTGATAAGCTCTAATGGCA-3' | rhamnose inducible expression |
| 89 | Fed7-nLuc-pSSR-fw | MB09_G5 | 5'-GAGTAGTGGAGGTTACCATATGGTGATAGCTGATCTAAATTTTCCTCC-3' | rhamnose inducible expression |
| 90 | Fed7-nLuc-pSSR-rev | MB09_G6 | 5'-CGGCAGCAGATCTCATCAAATGGGGATTGATTTGCGG-3' | rhamnose inducible expression |
| 91 | Fed8-nLuc-pSSR-fw | MB09_G7 | 5'-GAGTAGTGGAGGTTACCATATGCCCCATACCATTGTTACGG-3' | rhamnose inducible expression |
| 92 | Fed8-nLuc-pSSR-rev | MB09_G8 | 5'-CGGCAGCAGATCTCATGGCAGGGCTTTTCTGTAAATCAG-3' | rhamnose inducible expression |
| 93 | Fed9-nLuc-pSSR-fw | MB09_G9 | 5'-GAGTAGTGGAGGTTACCATATGAAGAAGCGTGTGACCCTG-3' | rhamnose inducible expression |
| 94 | Fed9-nLuc-pSSR-rev | MB09_H1 | 5'-CGGCAGCAGATCTCATGAAATTAGTGGCGATCGCCTGGACGGG-3' | rhamnose inducible expression |
| 95 | Fed10-nLuc-pSSR-fw | MB09_H2 | 5'-GAGTAGTGGAGGTTACCATATGGAAAACCCTGTCCCCATAAC-3' | rhamnose inducible expression |
| 96 | Fed10-nLuc-pSSR-rev | MB09_H3 | 5'-CGGCAGCAGATCTCATTAAATGCGAAAGAGGGTGGGTAAAAAAG-3' | rhamnose inducible expression |
| 97 | Fed11-nLuc-pSSR-fw | MB09_H4 | 5'-GAGTAGTGGAGGTTACCATATGACCATTACCTTTGTTAAAGAGCAGAAG-3' | rhamnose inducible expression |
| 98 | Fed11-nLuc-pSSR-rev | MB09_H5 | 5'-CGGCAGCAGATCTCATGCCTTTGGGTTTGGTGTTCAC-3' | rhamnose inducible expression |
| 99 | Fed12-nLuc-pSSR-fw | MB09_I9 | 5'-TAGAGTAGTGGAGGTTACCATATGGCAGTAACGATACACTTTTTGC-3' | rhamnose inducible expression |
| 100 | Fed12-nLuc-pSSR-rev | MB10_A1 | 5'-CGCTGCGGCAGCAGATCTCATCCAGGTGAGATCATCATAAAGATTAATTTCTAATTCC-3' | rhamnose inducible expression |
| 101 | Isib-nLuc-pSSR-fw | MB10_A8 | 5'-TAGAGTAGTGGAGGTTACCATATGACAAAAATTGGACTTTTTTACGGTACTCA-3' | rhamnose inducible expression |
| 102 | Isib-nLuc-pSSR-rev | MB10_A9 | 5'-CGCTGCGGCAGCAGATCTCATGGATTGCAAAATTGGTTTAATTTCACTTACC-3' | rhamnose inducible expression |
| 103 | pSSR_seq-fw | MB08_B9 | 5'-ATGGCCCATTTTCCTGTCAGTAACGAG-3' | mutant confirmation |
| 104 | pSSR_seq-rev | MB08_C1 | 5'-TTTGCTTCCAGATGTATGCTCTTCTGC-3' | mutant confirmation |
